# Benchmarking alternative polyadenylation detection in single-cell and spatial transcriptomes

**DOI:** 10.1101/2024.10.15.618405

**Authors:** Sida Li, Zixuan Wang, Yanshi Hu, Qingyang Ni, Cong Feng, Yueming Hu, Shilong Zhang, Ming Chen

## Abstract

**Background:** 3’-tag-based sequencing methods have become the predominant approach for single-cell and spatial transcriptomics, with some protocols proven effective in detecting alternative polyadenylation (APA). While numerous computational tools have been developed for APA detection from these sequencing data, the absence of comprehensive benchmarks and the diversity of sequencing protocols and tools make it challenging to select appropriate methods for APA analysis in these contexts.

**Results:** We systematically compared seven 3’-tag-based sequencing protocols and identified key peak features affecting APA detection performance. We developed a simulation pipeline that generates realistic datasets preserving protocol-specific characteristics. Using simulated and real data, we comprehensively assessed six computational tools for their ability to identify polyA sites, quantify polyA site expression, detect differentially expressed (DE) APA genes, filter sequencing artifacts, and their computational efficiency. We also investigated factors influencing APA detection. Our evaluation revealed that SCAPE and scAPAtrap generally outperformed other tools across various performance metrics and protocols.

**Conclusion:** Our systematic evaluation provides guidance for tool selection, experiment design, and future tool development in APA analysis for singlecell and spatial transcriptomics, paving the way for investigating APA in these contexts.

## 1 Introduction

Alternative polyadenylation (APA) is a widespread post-transcriptional regulatory mechanism. It generates mRNA isoforms with diverse 3’UTRs and terminal exons by utilizing different polyadenylation sites (pA sites) [1–3]. APA participates in numerous biological processes, such as disease progression, developmental regulation, and neural system modulation by affecting mRNA stability [4, 5], translation efficiency, subcellular localization [6, 7], and protein isoforms [8–10]. Bulk-level APA studies generally use specialized 3’-end sequencing technologies, which are similar to the 3’-tag-based RNA sequencing protocols commonly employed in single-cell and spatial transcriptomics. These protocols, such as 10X Chromium [11], Drop-seq, Microwell-seq [12], 10X Visium, Stereo-seq [13], Slide-seq V2 [14], and Spatial Transcriptomics (ST) [15], all generate sequence reads that are enriched at the 3’ ends of transcripts (Fig. 1b). Various tools have been developed to identify APA events from these 3’-tag-based sequencing data, such as scAPA [16], scAPAtrap [17], Sierra [18], SAPAS [19], scDaPars [20], SCAPTURE [21], MAAPER [22], SCAPE [23], and Infernape [24], enabling APA detection at single-cell and spatial resolutions. However, the absence of systematic benchmark and the diversity of sequencing protocols and APA detection tools make it challenging to select appropriate methods for APA detection in single-cell and spatial transcriptomics.

**Fig 1.**
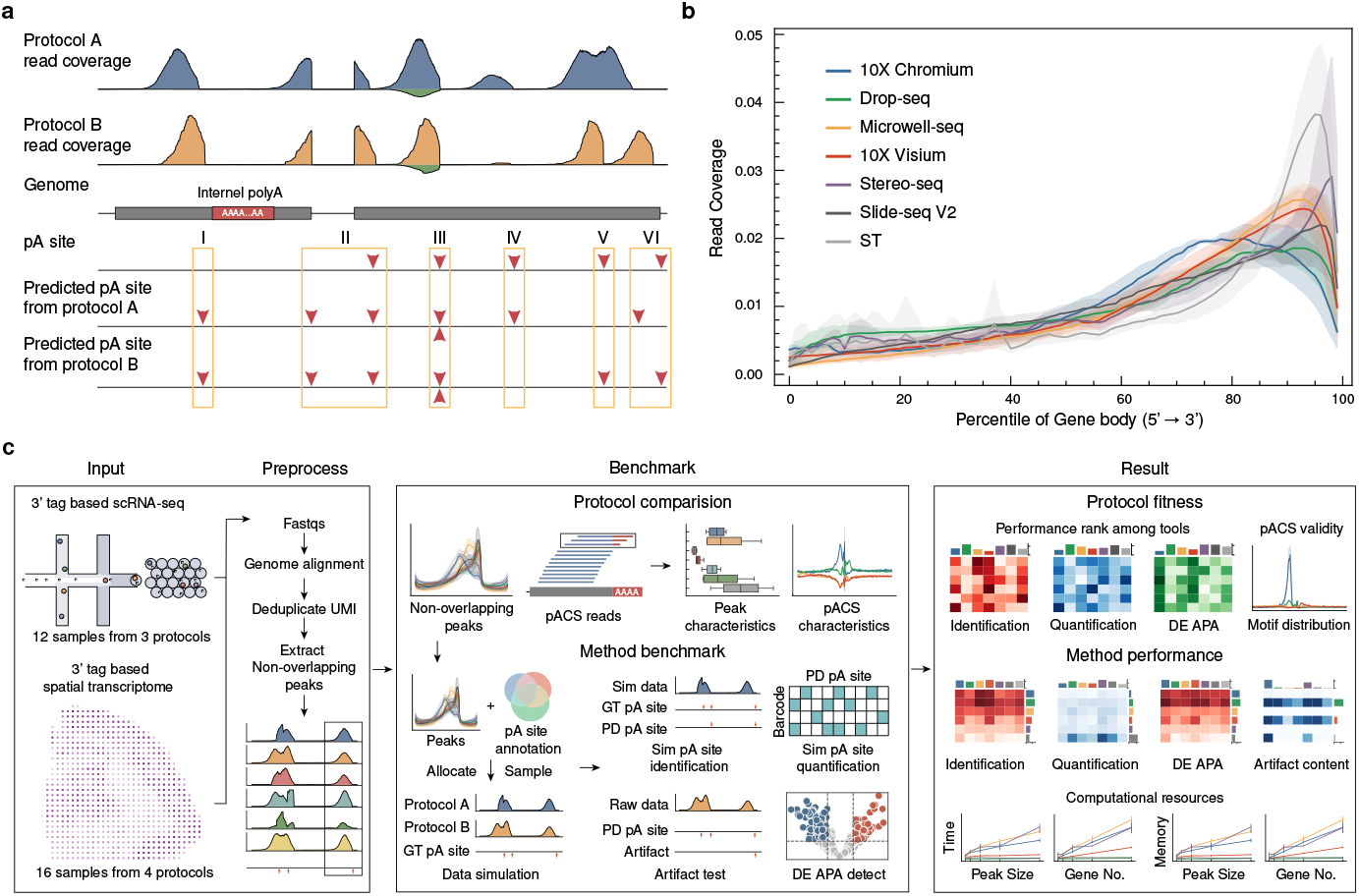
Challenges and feasibility of detecting APA events from 3’ enriched sequencing protocols and benchmarking workflow. (**a**) Challenges of detecting APA events from 3’ enriched sequencing protocols, (I) Internal oligo-dT priming; (II) Junction reads; (III) Antisense artifacts; (IV) Weak signal; (V) Peak overlap; (VI) Peak shift. (**b**) Read coverage by percentile of genebody across mainstream 3’-tag-based sequencing protocols. (**c**) Schematic overview of the benchmarking workflow. We evaluated the performance of 7 mainstream single-cell/spatial transcriptome sequencing protocols and 6 APA detection tools. For protocols, we assessed their suitability for APA event detection by examining peak characteristics and evaluating their performance in detecting APA events. For tools, we systematically evaluated their performance on simulated and real datasets, assessing their ability to identify polyA sites, quantify polyA site expression, detect differentially expressed (DE) APA genes, filter sequencing artifacts, and their computational efficiency.

Detecting APA events from 3’-tag-based RNA-seq data typically involves a four-step workflow: (i) identifying pA sites through peak calling; (ii) filter pA sites based on rules [16–19], statistical modeling [22, 23], or deep learning models [21]; (iii) quantifying pA site usage via rules [16, 17, 19, 23] or statistical modeling approaches [22–24]; (iv) Identify significant APA events based on statistical tests and thresholds of metrics that quantify APA usage. However, several challenges can complicate APA detection, including (i) peak overlap; (ii) peak shift; (iii) weak signals; and (iv) sequencing artifacts, such as junction reads, internal oligo dT, and antisense reads (Fig. 1a). These challenges vary in extent and impact across different sequencing protocols and samples, potentially leading to variable performance of APA detection tools. Current tool evaluations struggle to assess real-world performance due to the difficulty in accurately modeling peak characteristics in real data. Previous evaluations often rely on simulated data generated from probabilistic mixture model [23] or make comparisons on real data using pA site annotations as a substitute for the ground truth [17, 24, 25]. Both approaches have limitations that can hinder the accurate assessment of tool performance.

In this study, we systematically identified and compared data characteristics that could affect APA detection performance across seven 3’-tag-based sequencing protocols. These protocols cover a wide range of commonly employed sequencing chemistries and library preparation methods used in single-cell and spatial transcriptomics studies. We also evaluated the performance of six computational tools in detecting APA events using both simulated and real data, considering key metrics such as identification capability, quantification accuracy, differential APA detection capability, sequencing artifact filtering, and computational efficiency (Fig. 1c). This work provides guidance for researchers in selecting appropriate sequencing protocol and computational methods. Furthermore, it offers insights for developers to optimize APA detection tools, promoting the advancement of APA research in single-cell and spatial transcriptomics.

## 2 Results

### 2.1 Benchmark framework for sequencing protocols and tools that detects APA events from single-cell and spatial transcriptome

Our benchmark includes performance evaluations of different sequencing protocols and tools that detect APA events from single-cell and spatial transcriptomes (Fig. 1c). For sequencing protocols, we selected 7 mainstream single-cell/spatial transcriptome sequencing protocols, including 10X Chromium, Dropseq, Microwell-seq, 10X Visium, Stereo-seq, Slide-seq V2, and SpatialTranscriptomics (ST). Each protocol included 4 samples covering 2-3 mouse tissue types (Additional File 2: Supplementary Table 1). Since APA event detection from 3’ enriched sequencing data relies on peak calling, the peak characteristics of different sequencing protocols can affect the performance of APA detection tools. To examine the peak characteristics of each sequencing protocol, we extracted nonoverlapping peaks as representation based on integrated pA site annotations (See Methods). We evaluated peak characteristics including peak position, peak shape, and peak consistency. Furthermore, 3’ enriched RNA-seq data contains a certain number of reads with polyA cleavage sites (pACS), which could potentially serve as a reliable source for pA site identification. Therefore, we also assessed the feasibility of using pACS reads for pA site identification in different sequencing protocols by analyzing pA-site-related motifs, ATGC content, and conservation.

For tools, we collected as many APA detection tools applicable to spatial and single-cell transcriptomes as possible and selected 6 of them (Additional File 3: Table S2) for systematic performance evaluation on simulated and real datasets. We proposed a method to generate simulated data with ground truth while preserving protocol-specific peak characteristics. Specifically, we randomly sampled pA sites from the integrated mouse pA site annotation and assigned the nonoverlapping peaks from different protocols to these positions, ensuring that each pA site was associated with one peak. If the number of peaks was insufficient, existing peaks were duplicated and reassigned. We then distributed these peaks to different barcodes based on a negative binomial distribution and split them into two groups to simulate differential APA usage between cell/spot types (see Methods).

We evaluated the tools’ ability to identify pA site and multi-pA-site TEs using accuracy, recall, and F1-score as evaluation metrics. To assess identification consistency, we used the Jaccard index at pA site level and multi-pA-site TE level across sample replicates as metrics. For quantification capability, we used the mean absolute percentage error (MAPE) between the tools’ quantification results on simulated data and the ground truth pA site expression. To test how the aforementioned performance ultimately affects the identification of genes with significant APA differences, we used the F1-score, accuracy, and recall of differentially expressed (DE) APA genes under various statistical tests and filter thresholds as evaluation metrics. Furthermore, we assessed the ability of the six tools to filter out sequencing artifacts in real datasets, focusing on two common types of artifacts: reads spanning splice junctions and internal oligo(dT) primed reads. We calculated the proportion of identified pA sites originating from these two types of sequencing artifacts to evaluate each tool’s effectiveness in handling such artifacts. Finally, we assessed the computational resources consumed by each tool.

### 2.2 Comparison of peak characteristics from different sequencing protocols

To investigate the effect of peak characteristics from different sequencing protocols on APA detection, we compared peak features of seven sequencing protocols. We extracted nonoverlapping pA sites with no other pA site within 500 bp upstream or downstream of the integrated pA site annotation (see Methods). For each sequencing protocol, we retrieved all reads within 500 bp upstream and downstream of these nonoverlapping pA sites, obtaining a collection of nonoverlapping peaks. As shown in Fig. 2b,the majority of reads across all sequencing protocols were concentrated within the [-400,0] region surrounding the pA site (position 0), with a significant upward trend at 500 bp upstream and 200 bp downstream. On the basis of this observation, we selected reads from [-400,20] region surrounding the pA site to represent the peaks and discarded peaks containing fewer than 50 reads. Stereo-seq and 10X Chromium have the greatest number of nonoverlapping peaks, which is commensurate with their high read counts, whereas ST has the lowest read count and number of nonoverlapping peaks.

**Fig 2.**
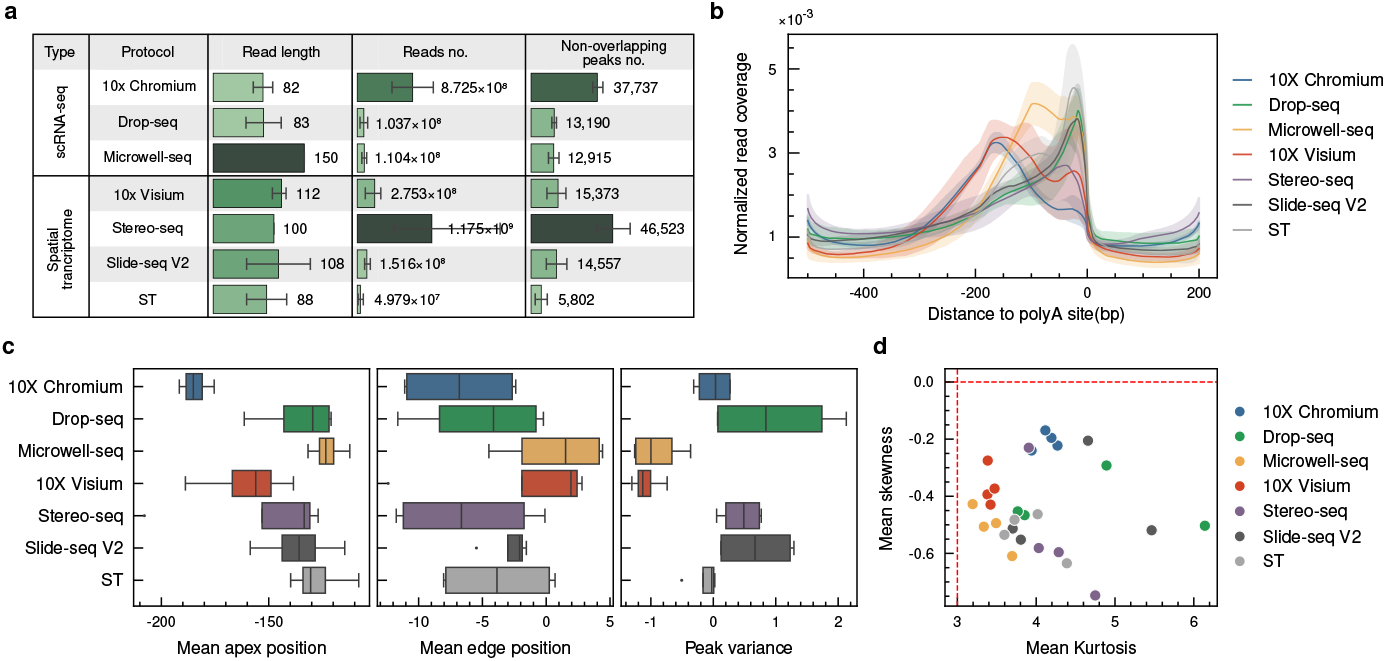
Peak characteristics of different sequencing protocols. (**a**) Summary of sequencing protocols. (**b**) Normalized read coverage around the pA site (position 0) for each sequencing protocol. The majority of reads were concentrated within the [-400,0] region. (**c**) Comparison of mean apex position, mean edge position, and peak variance across sequencing protocols. (**d**) Comparison of mean kurtosis and mean skewness across sequencing protocols.

We calculated five peak features for each sample: peak height, apex position (peak’s apex position relative to the pA site), edge position (peak’s upstream edge position relative to the pA site), kurtosis, and skewness (See Methods). For each feature, we computed the median and standard deviation (SD) across all peaks in a sample (Additional File 1: Fig. S1). The apex position varies among different sequencing protocols (Fig. 2c). 10X Chromium and 10X Visium presented the apex positions most distant from the pA site. Despite variations in the apex position, the edge positions were relatively consistent across protocols, with a maximum difference of less than 15 bp (Fig. 2c). We choose skewness and kurtosis as measures of peak shape. The majority of the peaks across all the protocols deviate from a normal distribution, exhibiting a negatively skewed and thin tail (Fig. 2d). Notably, the deviations were within an acceptable range for assuming normality (skewness in [-0.5, 0.5] and kurtosis in [1, 5]). Among the selected protocols, 10X Chromium produces peaks that most closely resemble a normal distribution. To further quantify the overall peak consistency across protocols, we introduced peak variance as a composite measure. The peak variance was calculated by averaging the z-scores of the standard deviations for apex position, edge position, kurtosis, and skewness. Microwell-seq and 10X Visium demonstrate the highest peak consistency, as evidenced by their low peak variance (Fig. 2c).

### 2.3 Feasibility of identifying pA site from reads with polyA cleavage sites

3’-tag-based sequencing technologies generate a small proportion of reads that contain polyA cleavage sites (pACSs), characterized by the presence of polyA sequences that fail to align to the reference genome at the 3’ end [17, 23]. We extracted reads containing pACS from seven sequencing protocols and noted that many putative pACSs had polyA (defined as a stretch of at least 10 continuous A with at most one mismatch) sequences in [-20,20] region (Additional File 1: Fig. S2b). Upon examining the alignment of these sites, we observed that alignment softwares resulted in errors in determining the end position of polyA sequences and aligned reads with long polyA to A-enriched regions of genome (Additional File 1: Fig. S2a). This suggests that this identification method may not completely eliminate the influence of internal oligo-dT priming. We then removed sites with polyA/polyT sequences around to avoid internal oligo-dT priming and potential antisense reads (see Methods). Filtered pACSs were then categorized into 3’UTR pACSs and non-3’UTR pACSs based on whether they could match 3’UTR annotations [26]. The number of pACS in different samples is shown in Fig. 3a. We analyzed the relationship between sequencing read length and pACS capture efficiency (defined as the ratio of pACS count to the total read count) and found that longer read lengths correlated with higher pACS capture efficiency (Spearman r=0.74, p=8.6e-11).

**Fig 3.**
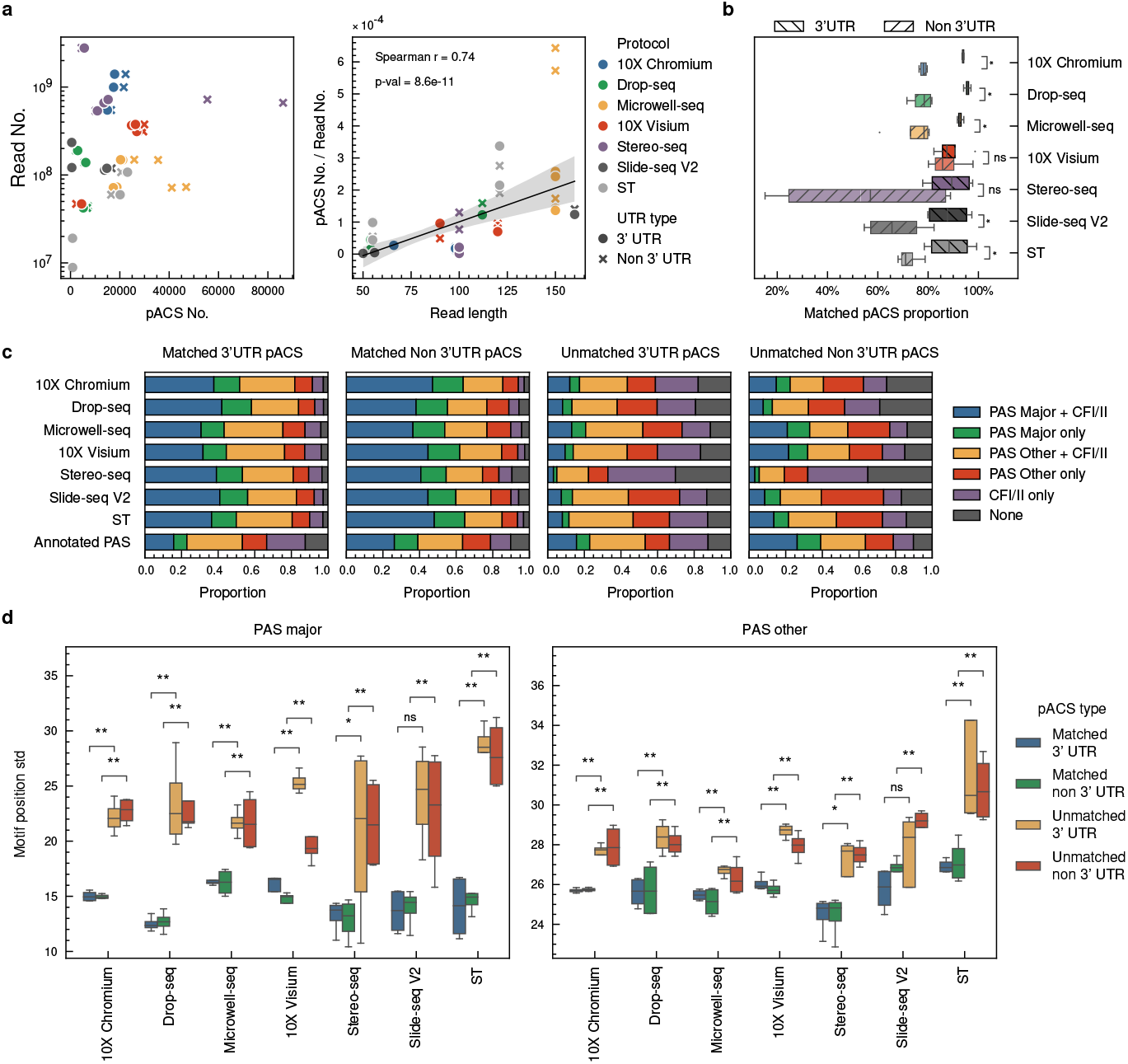
Feasibility of identifying pA sites from reads with polyA cleavage sites (pACS). (**a**) Number of pACS identified in different sequencing protocols (left) and the relationship between read length and pACS capture efficiency (right). (**b**) Proportion of pACS matching known pA site annotations in 3’UTR and non-3’UTR regions. (c) Proportion of pACS containing polyadenylation signal (PAS) motifs and cleavage factor (CF) binding sites. Motif definition (mark pACS as position 0): (i) PAS major, AATAAA in [-100,0] [3, 27]; (ii) PAS other, noncanonical PAS motifs in [-100,0] [27]; (iii) CFI, TGTA in [-100,0] [3]; (iv) CFII, TKTKTK in [0,100] [3]. (**d**) Standard deviation of PAS motif positions in different pACS categories.

To validate the reliability of pACS as pA sites, we matched pACS with a comprehensive mouse pA site annotation (Fig. 3b,see Methods). Nearly all samples had over 80% of their 3’UTR pACS matching known pA site annotations, with 10X Chromium, Drop-seq, and Microwell-seq exhibiting match rates greater than 90% for 3’UTR pACS. The match rates for non-3’UTR pACS were mostly ranging from 60% to 80%, which were significantly lower than those of 3’UTR pACS (Wilcoxon rank-sum test, p *<*0.05), except for Stereo-seq and Slide-seq V2. We observed an read enrichment upstream of pACS both at matched and unmatched pACS (Additional File 1: Fig. S3a,S3b). This enrichment pattern is consistent with the expected read coverage profile around genuine pA sites, where reads tend to accumulate upstream of the cleavage site.

We analyzed the binding sites of the cleavage and polyadenylation (CPA) mechanism around pACS. Functional pA sites typically include a polyadenylation signal (PAS), an upstream cleavage factor (CF) I binding site, and a downstream CFII binding site. We classified pACS based on the presence of specific motifs within designated windows (marking the pA site as position 0), including the PAS major motif (AATAAA in [-100,0]) [3, 27], PAS other motifs (noncanonical PAS motifs in [-100,0]) [27], CFI recognition motif (TGTA in [-100,0]) [3], and CFII recognition motif (TKTKTK in [0,100]) [3]. While the proportion of matched pACSs that contained PAS motifs was greater than that of the annotated pA sites, the proportion of PAS motifs in unmatched pACS was comparable to or slightly lower than that of annotated pA sites (Fig. 3c). Despite this, unmatched pACS still exhibited PAS enrichment upstream (Additional File 1: Fig. S5a), suggesting that at least some of these sites might be genuine pA sites. However, the PAS-motif-containing proportion in unmatched pACS from Stereo-seq was significantly lower than that in other protocols, indicating potential protocol-specific artifacts. To further assess the validity of unmatched pACS, we investigated their PAS motif distribution patterns and composition. Non-3’UTR pACS retained a clear upstream PAS motif distribution pattern, similar to that of annotated pA sites (Additional File 1: Fig. S4a). For 3’UTR pACS, while Microwell-seq, 10X Chromium, and Drop-seq maintained a discernible upstream distribution pattern, other sequencing protocols showed weaker enrichment (Additional File 1: Fig. S5a). Interestingly, the proportion of noncanonical PAS motifs was higher than that of canonical PAS motifs in unmatched pACS, which differs from the composition of annotated pA sites(Fig. 3c). Moreover, the PAS motif locations in unmatched pACS were more dispersed (Fig. 3d and Additional File 1: Fig. S5a). The more dispersed PAS motif locations and the higher proportion of noncanonical PAS motifs in unmatched pACS were consistent with the characteristics of minor pA sites, which often exhibit lower expression and higher cleavage site heterogeneity [28, 29].

We also computed the conservation around pACS (Additional File 1: Fig. S4c,S5c, see Methods). For 3’UTR pACS, we observed an enhanced conservation region upstream of the sites. In contrast, the conservation enhancement upstream of non-3’UTR pACS was weaker. When comparing these patterns to annotated pA sites, we found that the conservation patterns of pACS were consistent with the annotation, despite whether they matched the annotation, which further supports the reliability of pACS as genuine pA sites.

The base composition pattern surrounding pACS provides further evidence for the credibility of pA site identification. We focused on unmatched pACS and observed polyA enrichment upstream of pACS in 10X Chromium, Drop-seq, Microwell-seq, 10X Visium, and Slide-seq V2 (Additional File 1: Fig. S3c). This A-rich sequence context has been reported to be a primitive form of the PAS signal, which is more frequently found in younger, less conserved polyadenylation sites [30]. The presence of this A-rich sequence context upstream of unmatched pACS suggests that these sites might be evolutionarily young, undiscovered minor pA sites. However, Stereo-seq and ST did not exhibit corresponding patterns (Additional File 1: Fig. S5b). Notably, Stereo-seq exhibited a repeat-like pattern in the [-90,-70] region in both matched and unmatched pACS (Additional File 1: Fig. S4b, S5b), indicating a protocol-specific artifact. This artifact is likely caused by the rolling circle replication method employed by Stereo-seq.

In conclusion, except for Stereo-seq and ST, the pACS reads generated by other protocols can be used to identify pA sites after appropriate filtering. Samples with longer read lengths have higher pACS capture efficiency, providing an advantage in pACS identification.

### 2.4 Overall performance of APA detection tools

We evaluated the performance of six tools on 252 simulated datasets, which were generated based on the characteristics of seven sequencing protocols. For pA site identification, predicted pA sites within a 200-nt window centered on the ground truth pA site were considered valid, consistent with our data simulation criteria (see Methods).

To assess the ability of identifying multi-pA-site genes, we matched pA sites with terminal exons (TEs) and assigned the matched pA sites to the corresponding TEs. The tool’s ability to discover APA genes was evaluated based on its ability to predict multipA-site TEs. We found that the ability to identify pA sites and the ability to discover APA genes was highly correlated (Spearman r = 0.906, p *<*0.001). scAPAtrap and SCAPE outperformed other tools in identifying pA sites and discovering APA genes (Fig. 4a, 4b, and Additional File 1: Fig. S7). We also examined the consistency of pA site and multi-pA-site TE identification via the Jaccard index between replicate samples under the same ground truth and protocol. SCAPE, scAPA, and scAPAtrap showed good consistency in identifying pA sites and multi-pA site TEs (Fig. 4c, 4d). Furthermore, we investigated the ability of the tools to quantify pA site expression. We evaluated the mean absolute percentage error (MAPE) between the PAS expression matrix generated by the tools and the ground truth expression matrix at both barcode and group levels. Only matched pA sites were included in the calculation. If a predicted pA site matched multiple ground truths, it was assigned to the nearest one. If a ground truth matched multiple predicted pA sites, the sum of the expression values of the predicted pA sites was considered as the predicted expression value of that ground truth pA site. We found that the errors at the barcode level were generally small, with an average level below 0.1 except for Sierra, but these errors accumulated to a relatively large extent at the group level, reaching 0.15-0.3 except for Sierra (Fig. 4e, 4f). SCAPE outperformed the other tools in quantifying pA site expression.

**Fig 4.**
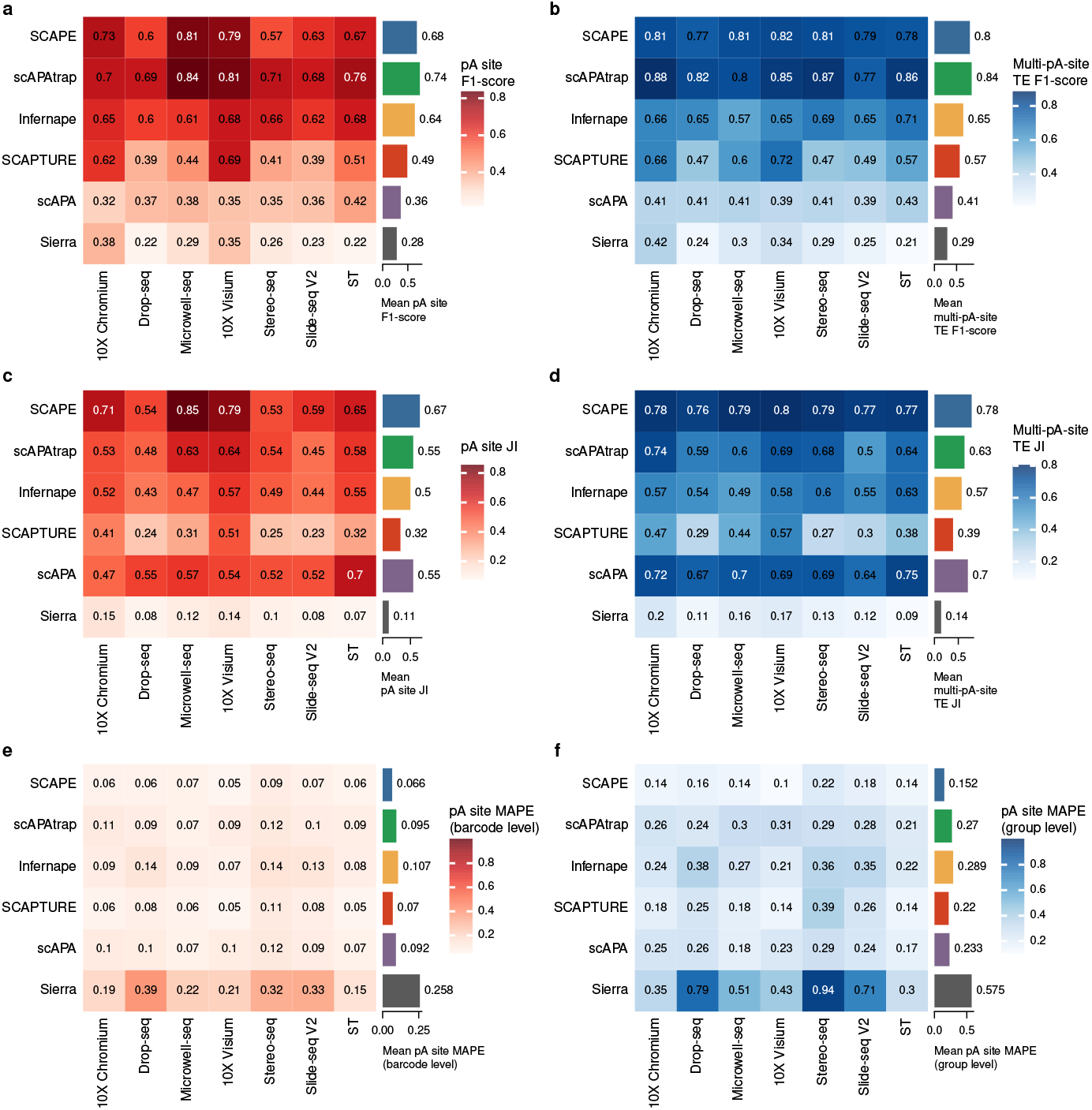
pA site identification and quantification performance of APA detection tools on simulated datasets. (**a**,**b**) Mean F1-score for (**a**) pA site and (**b**) multi-pA-site TE identification across different sequencing protocols. (**c**,**d**) Mean Jaccard index for (**c**) pA site and (**d**) multi-pA-site TE identification between replicate simulated samples under the same ground truth and protocol. (**e**,**f**) Mean absolute percentage error (MAPE) for pA site expression quantification at the (**e**) barcode and (**f**) group level across different sequencing protocols.

To assess the tools’ ability to identify differentially expressed (DE) APA genes in real-world scenarios, we selected combinations of three statistical tests and six filtering thresholds to screen for significant differences between two barcode groups in both the ground truth and the predicted results. The three statistical tests included the following: (i) Wilcoxon rank-sum test on one of the metrics that quantified APA usage, including percentage of distal poly(A) site usage index (PDUI), percentage of proximal poly(A) site usage index (PPUI), rank-weighted poly(A) site usage index (RWUI), or distance-weighted poly(A) site usage index (DWUI), between the two groups, with an FDR-corrected p-value *<*0.05 [16]; (ii) Fisher’s exact test, with an FDR-corrected p-value *<*0.05 [17, 24]; and (iii) DEXSeq [31] differential test, which randomly assigns two groups of barcodes into six pseudobulk subgroups, with any pA site in a TE group having a corrected p-value *<*0.05 [18, 23]. The six filtering thresholds included PDUI_diff_ *>*0.2 [20], PPUI_diff_ *>*0.2 [16], RWUI_diff_ *>*0.1 [32], DWUI_diff_ *>*0.1 [32], MPRO (maximum difference in proportion change) *>*0.2 [24], and DEXSeq log2FC *>*0.5 [18]. The precision, recall, and F1 score calculated from the filtered DE APA TEs were used as evaluation metrics. For each tool, we selected the filtering combination with the best performance as its representative. Compared with the other tools, scAPAtrap and SCAPE showed superior overall performance in identifying differential APA genes (Fig. 5a). Except for Infernape and scAPA, all the other tools achieved a prediction precision higher than 0.9 for all datasets (Fig. 5b). To further investigate the effect of pA site identification, multi-pA-site TE identification, and pA site quantification on the final performance of DE APA gene identification, we calculated the Spearman partial correlation between the performance metrics of these components and the performance metrics of DE APA gene identification. We found that the recall of multi-pA-site TE identification was most strongly correlated with the DE APA F1-score (Fig. 5d). The partial correlation analysis of the recall and precision of DE APA gene identification revealed that the recall of multi-pA-site TE identification affected the overall performance of DE APA gene identification by influencing the recall of DE APA (Fig. 5e, 5f). This finding was consistent with the high DE APA precision achieved by most tools (Fig. 5b). We hypothesized that factors such as pA site overlap, weak signals, and pA site shifting could influence the recall rate of multipA-site TE identification (Fig. 1a). To investigate this, we evaluated the performance of APA detection tools for TEs with varying pA site gaps and pA site read counts, and observed how the recall rate changed after extending the TE by different lengths (Additional File 1: Fig. S8). For each multi-pA-site TE, we ordered its actual pA sites based on their genomic positions and identified the pair of adjacent pA sites with the largest gap. We then used the gap size and the lower read count within this pA site pair as the representative features for the TE. Interestingly, all tools showed a performance drop for smaller pA site gaps, highlighting the substantial negative impact of pA site overlap on multi-pA-site TE recall. With the exception of Sierra, the recall rates of the other tools did not exhibit a notable decrease when the minimum pA site read count was reduced. We found that extending the TE by approximately 100 bp led to a slight improvement in recall rate across all tools, indicating that pA site shifting is a common phenomenon with a considerable influence. In essence, the primary limiting factor in identifying differentially expressed APA genes was the low recall rate of multi-pA-site TE identification, which was influenced by peak overlap and peak shift. We also tested the ability of the six tools to filter out sequencing artifacts, including junction-related pA site and internal-oligo-dT-related pA site, on the raw sequencing data. SCAPE, Infernape, and scAPA filtered sequencing artifacts, whereas scAPAtrap, SCAPTURE, and Sierra did not (Additional File 1: Fig. S6a, S6b).

**Fig 5.**
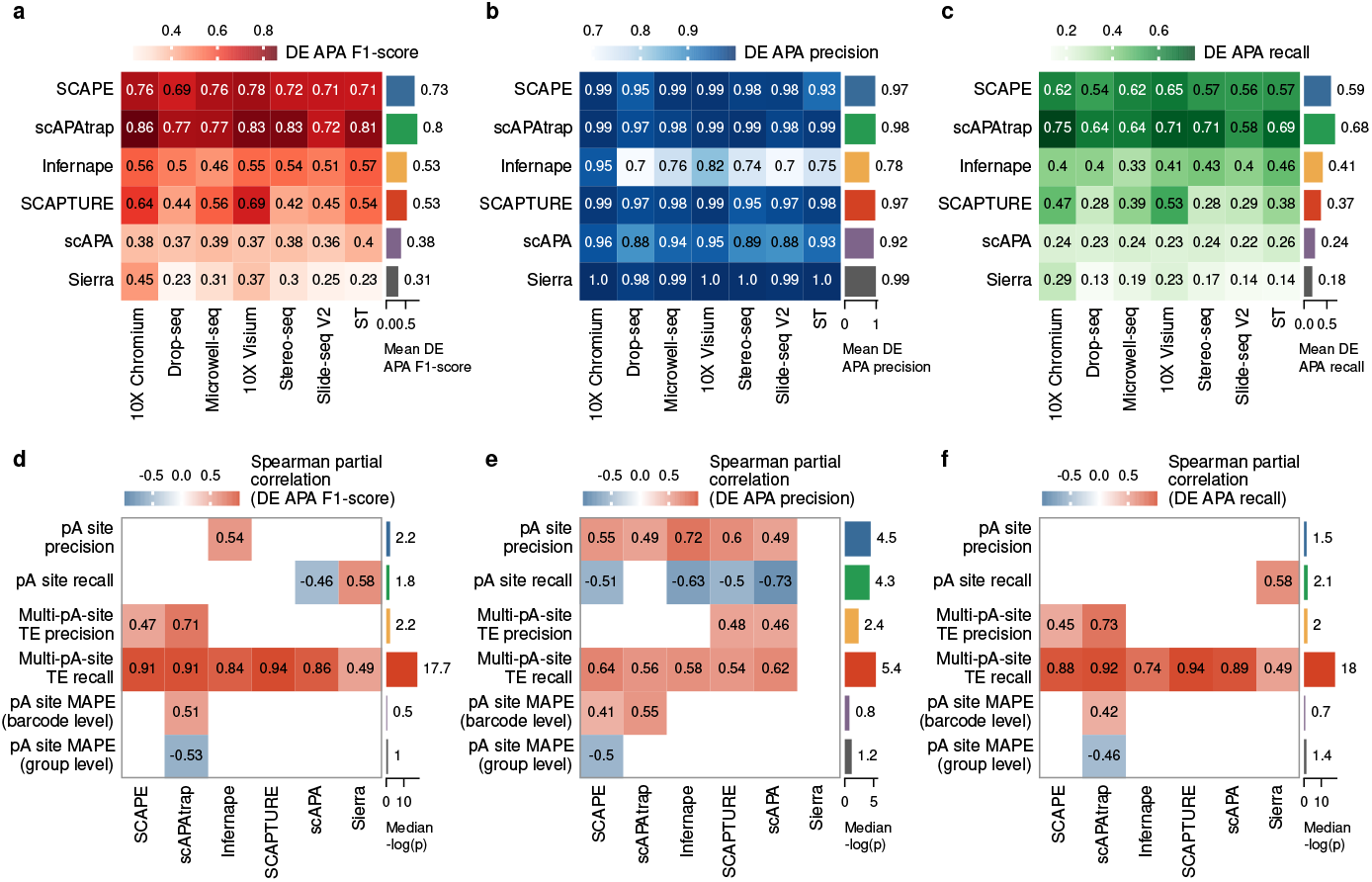
Performance evaluation of APA detection tools in identifying differentially expressed APA genes on simulated datasets. (**a**) Mean F1-score, (**b**) mean precision, and (**c**) mean recall for DE APA gene identification across different sequencing protocols. For each tool, the filtering criteria with the greatest F1-score is selected as its representative. (**d-f**) Spearman partial correlation between the performance metrics of pA site identification, multi-pA-site TE identification, pA site quantification, and the (**d**) F1-score, (**e**) precision, and (**f**) recall of DE APA gene identification. Correlations with p-value *>*0.05 were considered insignificant and removed from the heatmaps, with the corresponding cells left blank. The median −log(p) values represent the median of the negative logarithm of the p-values for the corresponding Spearman partial correlations across all datasets.

### 2.5 Effects of sequencing protocols and filtering criteria on APA detection

In addition to comparing the performance of different tools, we also investigated the differences in APA detection performance using simulated datasets generated from various sequencing protocols. We ranked the performance metrics of each tool on simulated datasets derived from the same ground truth but with protocol-specific characteristics and then calculated the average rank for each protocol (Fig. 6a). Although the tools exhibited varying preferences for different protocols, 10X Visium and 10X Chromium consistently outperformed other protocols across all the metrics, closely followed by Microwell-seq and ST. To identify protocol features that influence APA detection performance, we calculated the Spearman partial correlation coefficients between protocol peak characteristics and the performance metrics of each tool (Fig. 6b). The apex position SD had a significant negative effect on APA detection performance across multiple tools, particularly in terms of pA site expression quantification. This finding explains the relatively better performance of 10X Visium, 10X Chromium, Microwell-seq, and ST in APA detection, as they have lower apex position SD (Additional file 1: Fig. S1).

**Fig 6.**
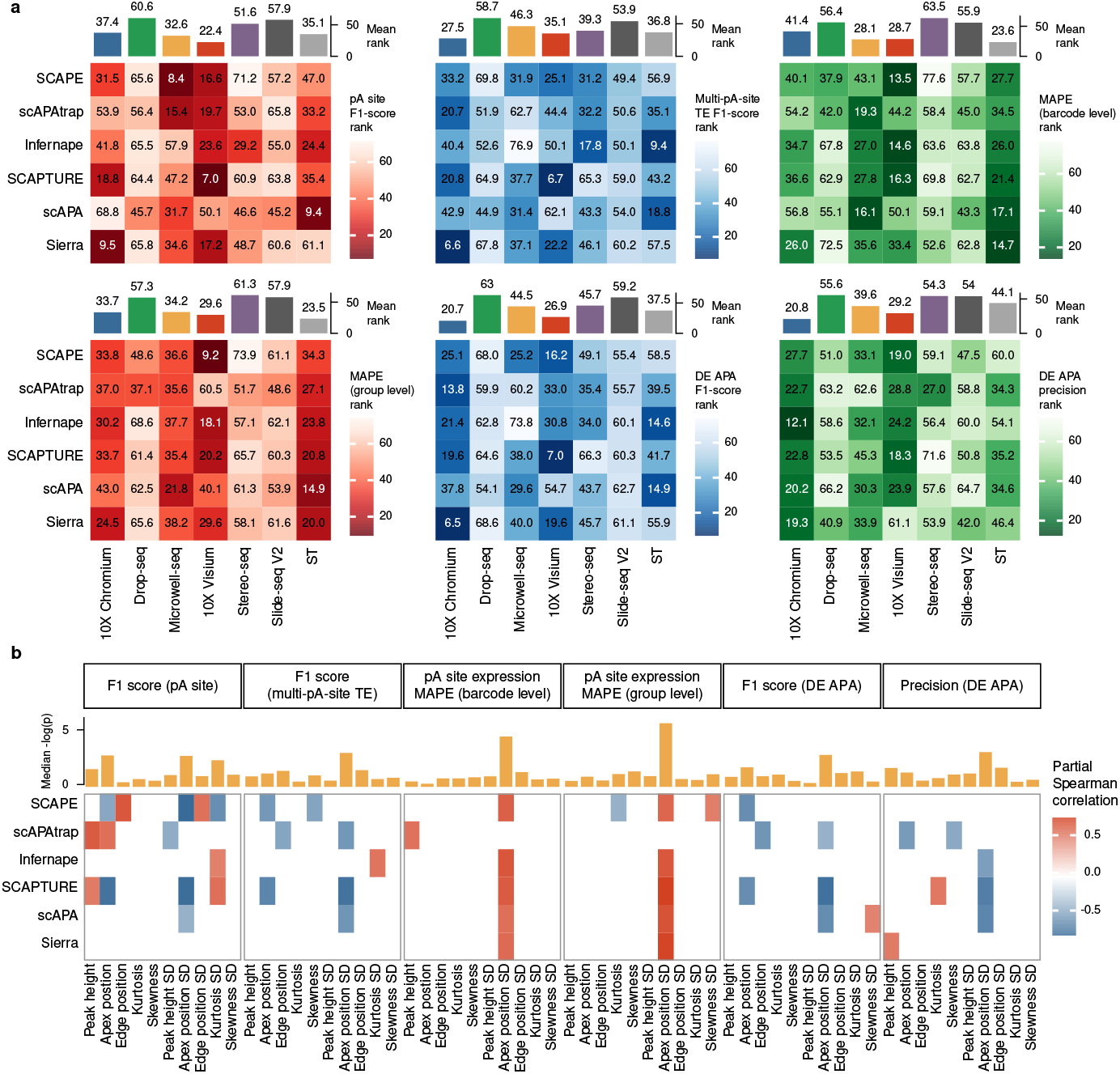
Effect of sequencing protocols on APA detection performance. (**a**) Heatmap showing the average performance rank of each sequencing protocol across different APA detection metrics. The metrics include F1-score for differentially expressed (DE) APA gene identification, precision for DE APA gene identification, mean absolute percentage error (MAPE) at the group level for pA site expression quantification, MAPE at the barcode level for pA site expression quantification, F1-score for pA site identification, and F1-score for multi-pA-site TE identification. Lower ranks indicate better performance. (**b**) Heatmap showing the Spearman partial correlation coefficients between protocol peak characteristics and the performance metrics of each tool. Correlations with p-value *>*0.05 were considered insignificant and removed from the heatmaps, with the corresponding cells left blank. The median −log(p) values represent the median of the negative logarithm of the p-values for the corresponding Spearman partial correlations across all datasets.

Next, we investigated the effect of filtering criteria on the identification of differentially expressed (DE) APA genes. We computed the F1-score and precision performance rankings for various combinations of statistical tests and filtering strategies within the same tool-sample combination. Under commonly used filtering thresholds, the combination of DEXSeq test/Fisher’s exact test and the DEXSeq log2FC filtering strategy demonstrated the best performance rankings. The F1-score rankings were similar across different statistical tests (Fig. 7a), but Fisher’s exact test and DEXSeq test significantly outperformed the Wilcoxon rank-sum test in terms of precision (Fig. 7b). The differences between filtering strategies were more pronounced than those between statistical tests, with DEXSeq log2FC exhibiting the best F1-score ranking (Fig. 7a), followed by RWUI_diff_ and MPRO. However, MPRO showed the highest precision ranking (Fig. 7b). Since specific threshold values may also affect performance, we expanded the threshold range and calculated the average performance for each threshold and strategy (Additional file 1: Fig. S8). Overall, the recall of DE APA gene identification decreased with increasing thresholds, while precision initially increased but then decreased. Within the expanded range, DEXSeq log2FC, RWUI_diff_, and MPRO still outperformed the other filtering strategies. Lowering the filtering thresholds can improve the robustness of the filtering process against the identification and quantification biases introduced by APA detection tools, but may result in biologically ambiguous filtering results.

**Fig 7.**
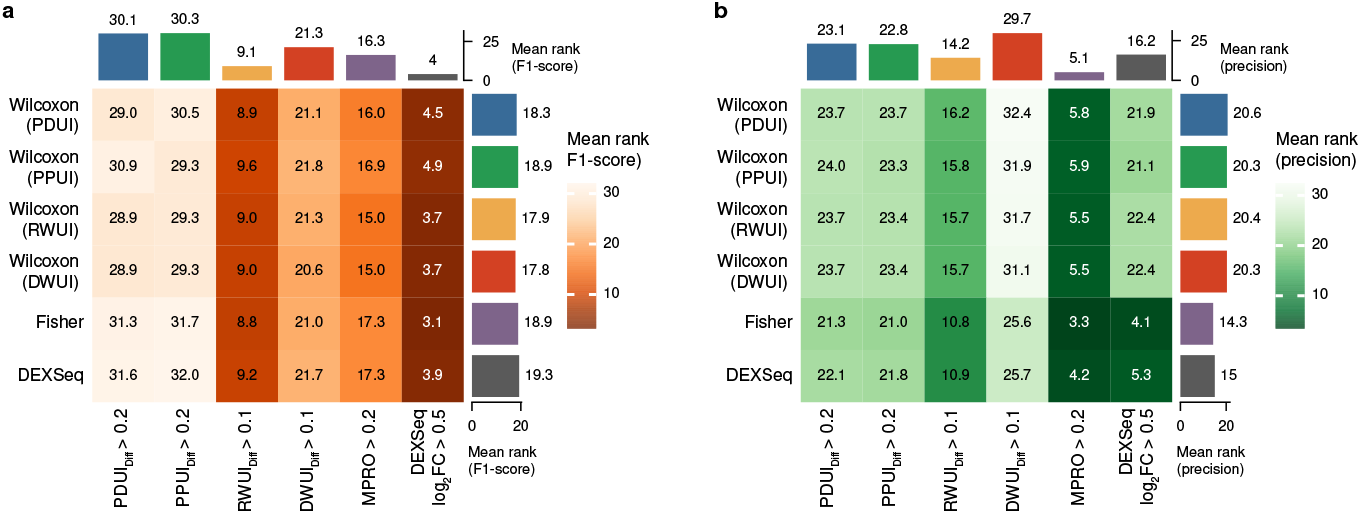
Effect of statistical tests and filtering strategies on the identification of differentially expressed APA genes. (**a**) Heatmap showing the average F1-score rank of different combinations of statistical tests and filtering strategies within the same tool-sample combination. Lower ranks indicate better performance.(**b**) Heatmap showing the average precision rank of different combinations of statistical tests and filtering strategies within the same tool-sample combination. Lower ranks indicate better performance.

### 2.6 Computational resource consumption of APA detection tools

We benchmarked the computational resource consumption of all APA detection tools under four distinct experimental conditions: (i) varying peak size (number of reads contained in a peak); (ii) varying gene number; (iii) varying barcode number; and (iv) varying read length. Each tool was configured uniformly to utilize eight processing cores, constrained by the `--cpus=8` option in Singularity. In terms of computation time, scAPAtrap significantly outperformed the other methods, closely followed by Sierra and SCAPTURE (Fig. 8a). For peak memory usage, SCAPTURE, SCAPE, and scAPAtrap greatly outperformed the other methods (Fig. 8b). Interestingly, although all tools claimed to support parallel computing, only SCAPE achieved full parallelization. The mean CPU load of scAPA increased significantly with increasing peak size and gene number, while those of scAPAtrap and SCAPTURE were closer to single-threaded computation (Fig. 8c).

**Fig 8.**
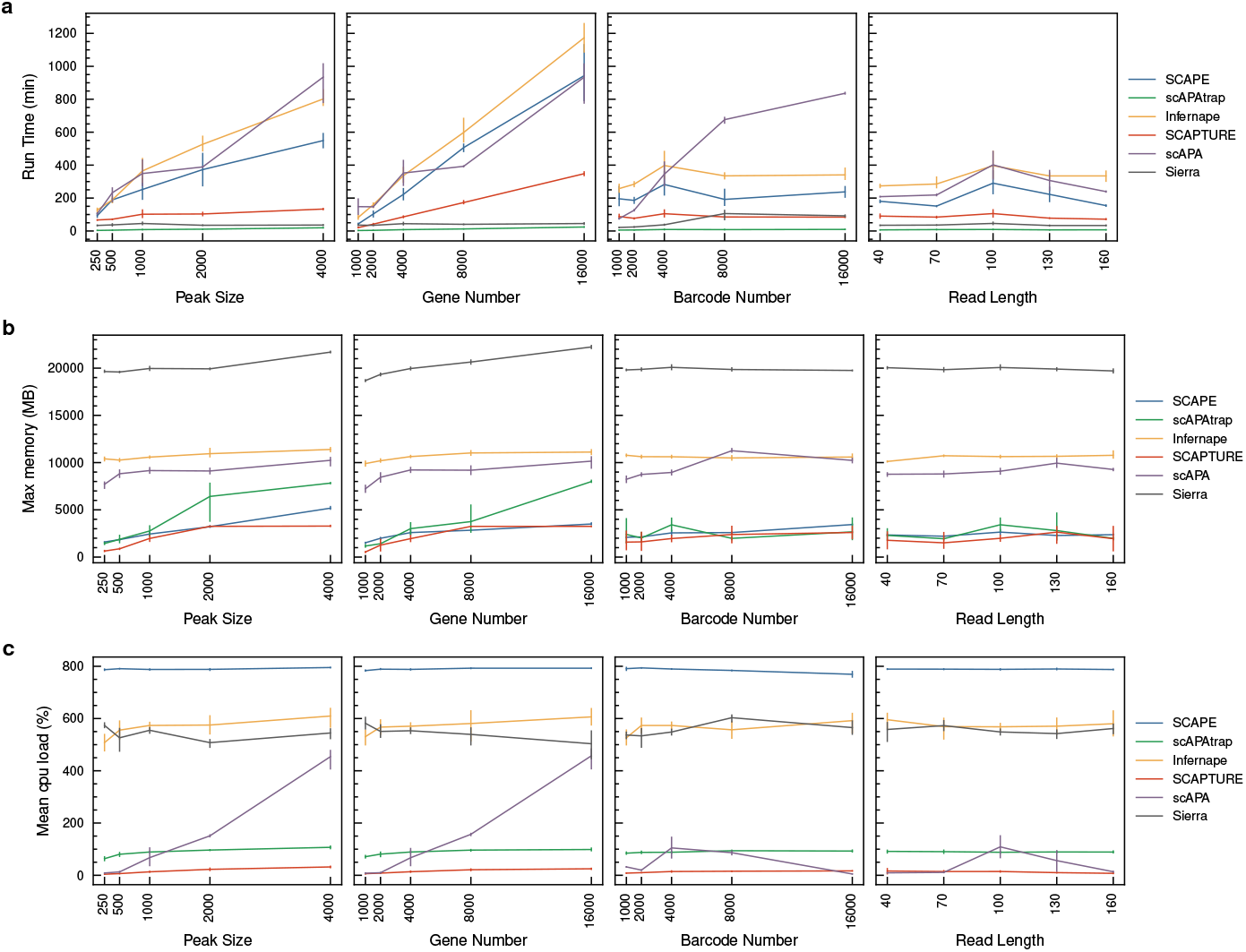
Computational resource consumption of APA detection tools under varying experimental conditions. (**a**) Run time (in minutes) of each tool under different peak sizes, gene numbers, barcode numbers, and read lengths. (**b**) Peak memory usage (in MB) of each tool under different experimental conditions. (**c**) Mean CPU load (in percentage) of each tool under different experimental conditions.

With respect to the effect of data parameters on computation, increasing the peak size and gene number significantly increased the computation time for all tools except scAPAtrap and Sierra, and slightly increased the peak memory consumption of all tools. This can be attributed to the subsequent filtering and statistical modeling of peaks performed by these tools, in which larger peak sizes and greater gene numbers significantly increase the postprocessing time but have a relatively minor effect on the peak calling step. Increasing barcode number had a significant negative effect on the running time of scAPA, while its effect on the running time of other tools was relatively minor. Read length had virtually no effect on the computational resource consumption of the tools.

## 3 Discussion

In this study, we established a comprehensive benchmarking framework to evaluate the performance of different sequencing protocols and tools in detecting APA events in single-cell and spatial transcriptome data. Our results revealed several key factors, such as the recall rate of multi-pA-site TE identification, peak characteristics of sequencing protocols, and the choice of filtering criteria, that significantly influence the performance of APA event detection. These findings provide important insights for optimizing existing methods and developing new APA detection tools.

When assessing the performance of different tools, we found that the recall rate of multi-pA-site TE identification is the main limiting factor affecting the performance of differential APA gene identification. This result emphasizes the importance of improving the sensitivity of multi-pA-site TE detection. We discovered that this limiting factor is influenced by pA site overlap and pA site shifting, revealing the inadequacies of existing peak calling methods and the importance of accurately identifying pA sites. The ability to recognize pA sites can be enhanced by integrating existing pA site annotations [22–24], employing deep learning models that incorporate genome sequence features [21], and applying statistical modeling to peaks [23, 24], thereby strengthening multi-pA-site TE identification.

Our analysis also revealed that peak characteristics, especially the standard deviation of apex positions, significantly affect APA detection performance. These characteristics were sample-specific, and influenced by a combination of factors such as the sequencing protocol, specific sample, library preparation, and sequencing platform. This finding highlights the necessity of considering sample specificity when designing and optimizing APA detection tools. Future tool development can explore strategies for optimizing algorithm parameters or feature selection tailored to different samples to improve the robustness of APA event detection. Previous studies have proposed utilizing pACS reads as a valuable information source for pA site identification [17, 32, 33]. However, our results suggest that the reliability of pACS reads varies not only among different sequencing protocols but also across individual samples within the same protocol. While some samples generate pACS reads that can be informative for pA site identification, others may produce pACS reads that were less reliable or contain errors introduced by various factors, such as sequencing artifacts and misalignment. Therefore, before considering the potential integration of pACS reads into existing APA detection tools or developing new tools that incorporate pACS information, it is crucial to carefully evaluate the quality and reliability of pACS reads in each specific sample. When evaluating the effect of filtering criteria on DE APA gene identification, we found that the combination of Fisher’s exact test/DEXSeq test with the DEXSeq log2FC filtering strategy performed the best. In terms of precision, MPRO and DEXSeq log2FC, which compare changes in pA site proportions, outperformed strategies that calculate changes in the overall APA usage index of the entire gene. This can be explained by the latter being more sensitive to incorrect pA site identification. As DEXSeq is a computationally intensive algorithm, computational resource pressures may arise when identifying DE APA genes across multiple cell/spot types. We recommend independently implementing the DEXSeq approach for calculating the normalized log2FC and then combining it with Fisher’s exact test for filtering.

Despite the comprehensive benchmarking and valuable insights provided by our study, there are several limitations that warrant further discussion. First, the simulated datasets were constructed using nonoverlapping peaks derived from the original data. However, due to potential incompleteness in annotations, a small proportion of overlapping peaks may have inadvertently been included in the nonoverlapping peak set. Second, to streamline data simulation and facilitate performance quantification, we focused exclusively on the most prevalent APA events occurring within TEs. Consequently, intronic APA and TE switch APA events, as described in [1–3], were not encompassed within the scope of this study. Furthermore, as only a subset of TEs were sampled and sequencing artifacts could not be systematically introduced into the simulated data, the resulting datasets may not fully capture the diversity and noise characteristics inherent to real-world data. Another limitation of our study is the limited number of sequencing samples used, which may not fully represent the variability within each protocol. Additionally, peak characteristics are not solely determined by the sequencing protocol but are influenced by a combination of factors, including the specific sample, library preparation, and sequencing platform. To generate simulated data that more faithfully recapitulates the complexities of real data for evaluating the performance of APA identification tools, further methodological innovations are necessary.

## 4 Conclusion

In this study, we established a comprehensive benchmarking framework to evaluate the performance of different sequencing protocols and tools in detecting APA events in single-cell and spatial transcriptome data. Among the tools evaluated, SCAPE and scAPAtrap exhibited the best overall performance in detecting APA events across various sequencing protocols and sample types. Furthermore, our results demonstrate that the combination of Fisher’s exact test/DEXSeq test with the DEXSeq log2FC filtering strategy outperforms other filter criteria in identifying differentially expressed APA genes. Our findings provide valuable insights and guidance for researchers studying APA events in single-cell and spatial transcriptome data, and can inform the development of future APA detection tools that consider sample-specific factors and implement optimal analysis strategies. This work provides a solid foundation for advancing the accuracy and efficiency of APA event detection in single-cell and spatial transcriptomics.

## 5 Methods

### 5.1 Data preprocessing

- 10X Chromium: Fastq files were aligned to the Gencode vm25 reference genome using cellranger-6.0.2.
- Drop-seq: Fastq files were aligned to the Gencode vm25 reference genome using
- Dropseq tools from https://github.com/broadinstitute/Drop-seq as described in [34].
- Microwell: Fastq files were aligned to the Gencode vm25 reference genome using the pipeline from https://github.com/ggjlab/mca data analysis as described in [12].
- 10X Visium: Fastq files were aligned to the Gencode vm25 reference genome using spaceranger-3.0.1.
- Stereo-seq: Fastq files were aligned to the Gencode vm25 reference genome using SAW6.0 from https://github.com/STOmics/SAW.
- Slide-seq V2: Fastq files were aligned to the Gencode vm25 reference genome using the slideseq-tools from https://github.com/MacoskoLab/slideseq-tools as described in [14].
- Spatial Transcriptomics: Fastq files were aligned to the Gencode vm25 reference genome using the ST pipeline [35] from https://github.com/SpatialTranscriptomicsResearch/st pipeline as described in [15].

After alignment, Uniquely mapped reads were extracted from the aligned BAM files using `samtools view -h -F 256 -bS`. PCR duplicates were then removed using `umi_tools` dedup.

### 5.2 An integrated annotation of mouse pA sites

To generate a comprehensive pA site annotation for the mouse genome, we integrated data from three well-established pA site databases: Gencode M25 [36], PolyA_DB v3.2 [37], and PolyASite 2.0[38], following [39]. First, pA sites within ± 10 nt of another pA sites within the same database were collapsed by selecting the most downstream pA sites. Next, the following procedure was implemented to reduce redundancy between databases: (i) we collected pA sites from PolyASite 2.0, (ii) we added pA sites from PolyA_DB v3.2 not within ± 10 nt of the current pA sites set, and (iii) we added pA sites from Gencode M25 not within ± 10 nt of the current pA sites set. This method of sequential addition led to a total of 539,346 pA sites in our union set. To streamline the subsequent data simulation steps, each pA sites in the unified set was assigned to the terminal exons of Gencode M25 gene annotations based on their genomic coordinates. In cases where a pA sites matched multiple terminal exons, it was duplicated in the final annotation.

### 5.3 Nonoverlapping peak analysis

A nonoverlapping pA site set was obtained by filtering the integrated pA site annotations, selecting those without any other pA sites within 500 bp upstream or downstream. For each sequencing technology, reads from the [-500, 200] region relative to each nonoverlapping pA site were extracted from the corresponding BAM files. Based on the distribution of reads flanking the nonoverlapping pA site, the [-400, 20] region was chosen to define pA-site-associated peak candidates. Nonoverlapping peaks containing fewer than 50 reads were discarded. The extracted peaks were converted into read coverage profiles for the calculation of peak features. for *peak*_*j,k*_ in *sample*_*k*_:

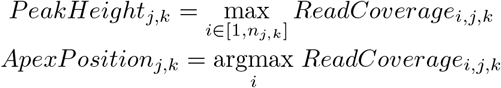

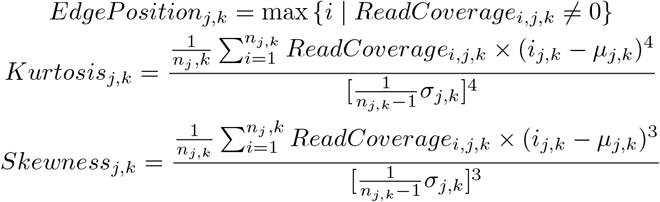

where:

- *ReadCoverage*_*i,j,k*_ denotes the read coverage at position *i* within the *peak*_*j*_ region in *sample*_*k*_.
- *n*_*j,k*_ is the number of positions within the *peak*_*j*_ region in *sample*_*k*_.
- *i*_*j,k*_ represents the position index within the *peak*_*j*_ region in *sample*_*k*_.
- *µ*_*j,k*_ is the weighted mean position of *peak*_*j*_ in *sample*_*k*_, calculated as:

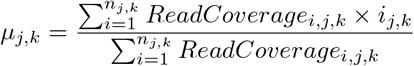
- *σ*_*j,k*_ is the weighted standard deviation of positions within the *peak*_*j*_ region in *sample*_*k*_, calculated as:

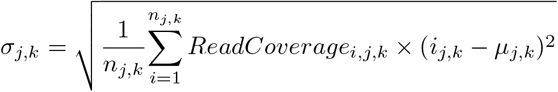

### 5.4 pACS analysis

We extracted reads containing pACS candidates from the aligned BAM files using the following criteria for the positive strand: (i) at least 10 nt of unaligned sequence at the 3’ end; (ii) allowing up to 2 non-A bases at the beginning of the unaligned sequence; (iii) at least 8 out of 10 consecutive bases are A, starting from the first A base in the unaligned sequence; (iv) the obtained pACS reads were merged using a 20 nt window to remove redundancy, retaining the pACS closest to the 3’ end as the representative of the merged group, and only pACSs supported by at least 2 reads were retained. For the negative strand, the criteria were adapted to consider the reverse complement of the sequences, with T bases instead of A bases. The resulting pACSs were searched for the presence of polyA/polyT sequences in a 40 nt window upstream and downstream using `seqkit grep`, with polyA/polyT defined as a stretch of 10 consecutive A/T allowing up to 1 mismatch. pACSs containing polyA/polyT sequences within this window were removed from subsequent analyses. We matched the pACSs to the integrated polyA site annotations using a 20 nt window and classified them as matched or unmatched based on whether they could be matched to the mouse pA site annotation. To filter for 3’UTR pACSs, we extracted 3’ UTR annotations from the mm39 UTR annotations obtained from UTRdb2.0 [26] and converted them to mm10 using UCSC liftOver. We then classified the pACSs as 3’UTR pACSs or Non-3’UTR pACSs based on whether they matched the 3’UTR annotations. For the base composition analysis, we calculated the ATGC content within a 400 nt window centered on the pACS site. For the conservation analysis of pACSs, we utilized the phastCons60way mm10 profile to extract a 200-nucleotide window centered on each pACS, excluding annotated coding sequence regions. We then calculated the mean conservation score at each position within the window by aggregating the conservation scores across all pACSs identified in each sequencing sample. For the pA site related motif analysis, we used the motif class from Biopython [40] to search for the corresponding motifs within a 200 nt window centered on the pACS site. The motifs searched included PAS major: canonical PAS hexamer (AATAAA) [3, 27]; PAS other: non-canonical PAS motifs (ATTAAA, AGTAAA, TATAAA, CATAAA, TTTAAA, AATACA, AATATA, GATAAA, AAGAAA, AATGAA, AATAGA, ACTAAA) [27]; CFI: CFI recognition motif (TGTA) [3]; CFII: CFII recognition motif (TKTKTK) [3]. For the classification of motifs contained in the pACS sites, we selected PAS major in the [-100,0] region, PAS other in the [-100,0] region, the CFI recognition motif in the [-50,0] region, and the CFII recognition motif in the [0,100] region.

### 5.5 Data simulation

To comprehensively evaluate the performance of the proposed method, we designed a simulation scheme to generate realistic 3’-tag-based RNA-seq data with known APA events, preserving peak characteristics from different sequencing protocol. The simulation process consists of the following steps:

#### 1. Peak extraction

We used the nonoverlapping peaks we extracted from each sample as described above, reads with the [-400, 20] region were chosen and peaks with fewer than 50 reads were dropped. CIGAR string and distance to pA site of each selected reads were recorded for read regeneration. Read counts for each *peak*_*j*_ are recorded as *c*_*j*_.

#### 2. Gene and terminal exon set definition

We define the following gene and terminal exon sets based on the number and differential usage of pA site:

- Single-pA-site gene set *G*_*single*_:

- This set contains all genes with only one pA site. Formally, if gene *g*_*j*_ has only one pA site, then *g*_*j*_ ∈ *G*_*single*_. This set is designed for tools like Infernape [24] that require single-pA-site genes for modeling.
- Multi-pA-site terminal exon set *T*_*multi*_:

- This set contains all terminal exons with multiple pA site, each with at most five pA site. Formally, if terminal exon *t*_*j*_ has multiple pA site, then *t*_*j*_ ∈ *T*_*multi*_. Adjacent pA sites are at least 200 bp away.
- *T*_*multi*_ can be divided into two subsets:

* Differential APA terminal exon set *T*_*diff*_ : This subset contains terminal exons with pA site showing differential usage between cell/spot types.
* Nondifferential APA terminal exon set *T*_*nondiff*_ : This subset contains terminal exons with pA site showing no significant differential usage between cell/spot types or conditions.

Gene set and terminal exon set were randomly sampled from the mouse integrated pA site annotation described above. If a selected TE contains more than five pA site, we will randomly choose *n* pA sites as representation, where *n*∼ *𝒰* { 2, 3, 4, 5}. For each simulated BAM file in this study, we sampled 1000 single-pA-site genes, 2000 differential APA terminal exons, and 2000 nondifferential APA terminal exons. This sampling process was repeated three times to generate three independent datasets, ensuring the robustness of the simulation.

The nonoverlapping peaks were randomly assigned to each pA site without replacement. If the number of peaks is insufficient to cover all pA site, existing peaks will be duplicated and reassigned to guarantee that each pA site was associated with one peak. The read count for the peak assigned to *pAsite*_*j,k*_ within gene (*g*_*j*_) or terminal exon (*t*_*j*_) was recorded as *c*_*j,k*_.

#### 3. Gene expression allocation

The total expression level of the gene (*g*_*j*_) or terminal exon (*t*_*j*_) with *n* pA site is calculated as:

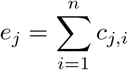

The relative usage proportion of each pA site is calculated as:

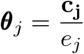

For each gene (*g*_*j*_) or terminal exon (*t*_*j*_), we generate an allocate vector **Δ**_*j*_ to allocate gene expression and simulate APA usage between cell/spot type *T*_1_ and *T*_2_. Each element Δ_*j,i*_ of **Δ**_*j*_ is sampled from a Beta distribution:

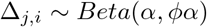

where *α* is a shape parameter and *ϕ* is a scaling factor uniformly sampled from the range [0.25, 4]. By restricting *ϕ* to this range, we ensure that the expected value of the Beta distribution falls between 0.2 and 0.8, allowing us to control the allocation of reads between the two cell/spot types. Additionally, we randomly swapped adjacent elements of **Δ**_*j*_ with a probability of 0.1 to introduce further randomness.

We then use **Δ**_*j*_ to allocate **c**_*j*_ to *T*_1_ and *T*_2_, obtaining **c**_*j,T*_1 and **c**_*j,T*_2, respectively:

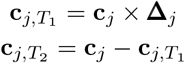

The expression levels 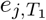 and 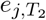 and the pA site usage proportion vectors 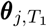 and 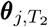 are calculated accordingly.

To ensure that the simulated pA site usage proportions exhibit the desired level of difference between the two cell/spot types, we perform a chi-square test on ***θ***_*j,T*_1 and ***θ***_*j,T*_2 for multi-pA site terminal exons. For terminal exons in the differential APA set *T*_*diff*_, we require a p-value *<*0.05 and a proportion difference (PD) *>*0.2. For those in the nondifferential set *T*_*nondiff*_, we require a p-value *>*0.1 and a PD *<*0.2. If these criteria are not met, we regenerate the allocate vector **Δ**_*j*_ and repeat the process until the desired level of difference or similarity is achieved.

#### 4. Expression simulation

For each cell/spot type *k* (*T*_1_ or *T*_2_) with *m* cell/spots, we simulated the gene expression levels and pA site usage proportions as follows:

(a) Determine the number of non-zero cell/spots *m*_*j,k,nonzero*_: For terminal exon *t*_*j*_ in cell/spot type *k*, we set the number of non-zero cell/spots as:

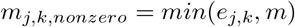

This ensures that the average expression of terminal exon *t*_*j*_ in expressing cell/spots is at least 1. The remaining cell/spots are assigned zero expression to simulate dropouts in single-cell RNA-seq data.
(b) Sample gene expression levels: For each non-zero cell/spot *i* in cell/spot type *k*, the expression level *e*_*j,k,i*_ of gene (*g*_*j*_) / terminal exon (*t*_*j*_) is sampled from a negative binomial distribution:

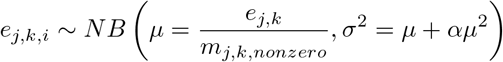

where *α* = 0.1 is used to adjust the dispersion of the distribution.
(c) Assign reads to pA site: Given the expression level *e*_*j,k,i*_ of terminal exon *t*_*j*_ in cell/spot *i* of cell/spot type *k*, we sample the read counts for each pA site using a multinomial distribution for multi-pA site terminal exons:

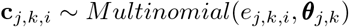

For single-pA site genes, we directly assign the expression level to the unique pA site:

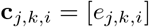

#### 5. Read generation

Finally, we generated reads by extracting sequences from the reference genome based on the pA site coordinates, read distance to pA site, and CIGAR string. We then assemble them into reads with simulated UMIs and barcodes utilizing previously assigned pA site counts for each cell/spot. For each real sample from different sequencing protocols, we generated three replicates using three independent pA site ground truth annotations to ensure the robustness of the simulation. Simulated reads were formatted and dumped as BAM file.

### 5.6 Settings for APA detection tools

- **SCAPE**: We followed the instructions on the SCAPE website: https://github.com/LuChenLab/SCAPE. Default parameters were used in the analysis.
- **scAPAtrap**: We followed the instructions on the scAPAtrap website: https://github.com/BMILAB/scAPAtrap. Default parameters were used in the analysis.
- **Infernape**: We followed the instructions on the Infernape website: https://github.com/kangbw702/Infernape. Default parameters were used in the analysis.
- **SCAPTURE**: We followed the instructions on the SCAPTURE website: https://github.com/kangbw702/Infernape. Default parameters were used in the analysis.
- **scAPA**: We followed the instructions on the scAPA website: https://github.com/ElkonLab/scAPA. Default parameters were used in the analysis. The mouse intron and UTR annotations used in scAPA are based on the mm9 genome version. We used UCSC liftOver to convert these annotations to the mm10 version.
- **Sierra**: We followed the instructions on the Sierra website: https://github.com/VCCRI/Sierra. Default parameters were used in the analysis.

scAPAtrap, scAPA, Sierra, SCAPTURE, and Infernape provide intervals of the peaks corresponding to predicted polyadenylation (pA) sites, but do not provide the exact locations of the pA sites. We used the end point of each interval (nearest to the poly(A) tail) as the predicted location of the pA site for these methods.

### 5.7 Benchmarking evaluation metrics

- **pA site identification evaluation**: Predicted pA sites were matched to ground truth pA sites using BEDTools window with a 200-nt window. *TP*_*pA*_ represents the number of predicted pA sites that matched ground truth pA sites, *FP*_*pA*_ represents the number of predicted pA sites that did not match any ground truth pA sites, and *FN*_*pA*_ represents the number of ground truth pA sites that were not matched by any predicted pA sites.

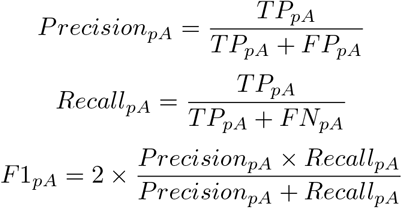

To assess the consistency of pA site identification, we calculated the Jaccard index between each pair of replicates within the same sample, ground truth, tool, and protocol combination, considering the sets of predicted pA sites that matched ground truth pA sites. The Jaccard index between two replicates *i* and *j* is defined as:

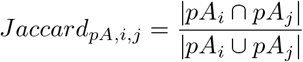

where *pA*_*i*_ and *pA*_*j*_ are the sets of predicted pA sites that matched ground truth pA sites in replicates *i* and *j*, respectively. For each combination of sample, ground truth, tool, and protocol with *n* replicates, the mean Jaccard index across all pairwise comparisons of the replicates was used to represent the consistency of pA site identification:

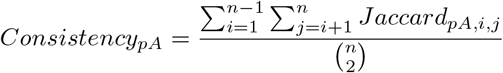
- **multi-pA-site TE identification evaluation**: Ground truth TEs were classified as multi-pA-site TEs if they contained more than one ground truth pA site. Predicted pA sites were matched to ground truth TEs using BEDTools intersect, and predicted TEs were classified as multi-pA-site TEs if they contained more than one predicted pA site. *TP*_*T E*_ represents the number of TEs that were classified as multipA-site TEs in both the ground truth and the predictions, *FP*_*T E*_ represents the number of TEs that were classified as multi-pA-site TEs in the predictions but not in the ground truth, and *FN*_*T E*_ represents the number of TEs that were classified as multi-pA-site TEs in the ground truth but not in the predictions.

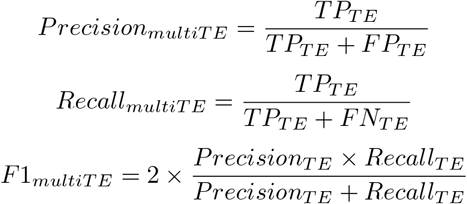

To assess the consistency of multi-pA-site TE identification, we calculated the Jaccard index between each pair of replicates within the same sample, ground truth, tool, and protocol combination, considering the sets of predicted multi-pA-site TEs. The Jaccard index between two replicates *i* and *j* is defined as:

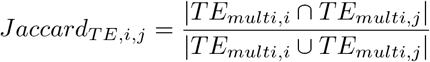

where *TE*_*multi,i*_ and *TE*_*multi,j*_ are the sets of predicted multi-pA-site TEs in replicates *i* and *j*, respectively. For each combination of sample, ground truth, tool, and protocol with *n* replicates, the mean Jaccard index across all pairwise comparisons of the replicates was used to represent the consistency of multi-pA-site TE identification:

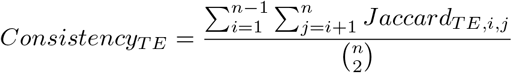
- **pA site quantification evaluation** We evaluated the accuracy of pA site quantification by calculating the mean absolute percentage error (MAPE) between the PAS expression matrix generated by the tools and the ground truth expression matrix at both barcode and group levels. Only matched pA sites were included in the calculation. If a predicted pA site matched multiple ground truth pA sites, it was assigned to the nearest site. If a ground truth pA site matched multiple predicted pA sites, the sum of the expression values of the predicted pA sites was considered as the predicted expression value of that ground truth pA site. The MAPE at the barcode level for a given sample, ground truth, tool, and protocol combination is defined as:

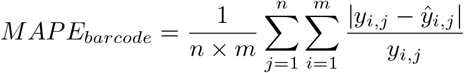

where *n* is the number of barcodes, *m* is the number of matched pA sites with non-zero ground truth expression values in the sample, *y*_*i,j*_ is the ground truth expression value of the pA site *i* in barcode *j*, and *ŷ*_*i,j*_ is the predicted expression value of the pA site *i* in barcode *j*. To avoid division by zero, pA sites with ground truth expression values equal to zero were excluded from the MAPE calculation. At the group level, barcodes are aggregated into predefined groups. The expression value of a pA site in a group is the sum of the expression values of the corresponding pA site across all barcodes in that group. The MAPE at the group level for a given sample, ground truth, tool, and protocol combination is defined as:

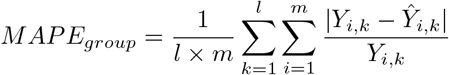

where *l* is the number of groups, *m* is the number of matched pA sites with non-zero ground truth expression values in the sample, *Y*_*i,k*_ =∑ _*j*∈*k*_ *y*_*i,j*_ is the ground truth expression value of the pA site *i* in group *k*, and *Ŷ*_*i,k*_ =∑ _*j*∈*k*_ *ŷ*_*i,j*_ is the predicted expression value of the pA site *i* in group *k*.
- **DE APA gene identification evaluation** To assess the tools’ ability to identify differentially expressed (DE) APA genes, we selected combinations of three statistical tests and six filtering thresholds to screen for significant differences between two barcode groups in both the ground truth and the predicted results.

The three statistical tests included:

- Wilcoxon rank-sum test on one of one of the four APA usage indices: percentage of distal poly(A) site usage index (PDUI), percentage of proximal poly(A) site usage index (PPUI), rank-weighted poly(A) site usage index (RWUI), or distance-weighted poly(A) site usage index (DWUI) between two groups, with an FDR-corrected p-value *<*0.05 [16];
- Fisher’s exact test, with an FDR-corrected p-value *<*0.05 [17, 24];
- DEXSeq [31] differential test, which randomly assigns two groups of barcodes into six pseudobulk subgroups, with any pA site in a TE group having a corrected p-value *<*0.05 [18, 23].

The six filtering thresholds included:

- PDUI_diff_ *>*0.2 between tested groups[20];
- PPUI_diff_ *>*0.2 between tested groups[16];
- RWUI_diff_ *>*0.1 between tested groups[32];
- DWUI_diff_ *>*0.1 between tested groups[32];
- MPRO (maximum difference in proportion change) between tested groups*>*0.2 [24];
- -Any pA site in a TE group having a DEXSeq corrected p-value *<*0.05 and log2FC *>*0.5 between tested groups[18].

For a gene containing *n* pA sites, we order the pA sites based on their genomic positions from upstream to downstream as (*p*_1_, *p*_2_, …*p*_*n*_). Considering the most upstream pA site as the origin, the positions of each pA site are denoted as (*d*_1_, *d*_2_…*d*_*n*_), and the expression levels of each pA site are represented as (*c*_1_, *c*_2_…*c*_*n*_). The following metrics are defined:

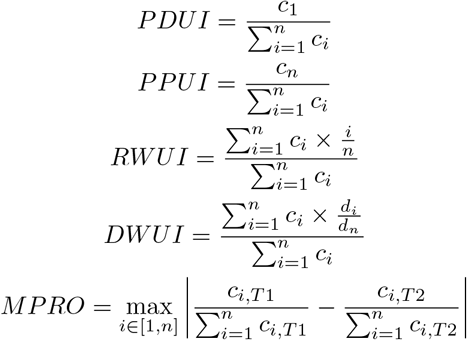

Ground truth DE APA genes were identified by applying the aforementioned statistical tests and filtering thresholds to the ground truth TEs. Similarly, predicted DE APA genes were identified by applying the same tests and thresholds to the predicted TEs. *TP*_*DEAP A*_ represents the number of genes that were classified as DE APA genes in both the ground truth and the predictions, *FP*_*DEAP A*_ represents the number of genes that were classified as DE APA genes in the predictions but not in the ground truth, and *FN*_*DEAP A*_ represents the number of genes that were classified as DE APA genes in the ground truth but not in the predictions. The precision, recall, and F1 score for DE APA gene identification are defined as follows:

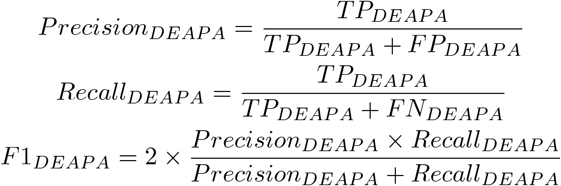

### 5.8 Computational resource benchmark

To evaluate the relationship between computational resource consumption and data characteristics for different tools, we generated a series of simulated datasets using data simulation pipeline described above, with specified parameters. The simulation process used peaks sampled from a normal distribution as input instead of peaks extracted from sequencing data. We set gene number=4000, peak size=1000, barcode number=4000, and read length=100 as the standard sample parameters. We then performed single-variable controlled simulations by varying gene number (1000, 2000, 4000, 8000, 16000), peak size (250, 500, 1000, 2000, 4000), barcode number (1000, 2000, 4000, 8000, 16000), and read length (40, 70, 100, 130, 160). The generated datasets were processed using tool benchmark pipeline described above, and the computational resource consumption was recorded using the benchmark function of Snakemake. The number of CPUs was limited to 8 by the --cpus=8 in Singularity, as some tools (such as scAPA) may exceed the specified CPU limit written in script. We conducted the performance tests of the six APA detection tools on a computer with two AMD Epyc 7763 CPUs (2.45 GHz, 256 MB L3 cache, 128 CPU cores in total) and 1 TB of memory (DDR4 2400 MHz).

## Supplementary information

## Declarations

- Ethics approval and consent to participate Not applicable.
- Consent for publication Not applicable.
- Availability of data and materials Code to reproduce all results and figures is available at Lycidas97/apabenchmark. All data and code are released under a GNU General Public License. Sequencing datasets used in this study are summarized in Additional file 2: Table S1 and are all publicly available at GSE121891 [41], GSE184708 [42], GSE117011 [43], GSE112393 [44], GSE153562 [45], GSE120374 [46], GSE169021 [47], PRJNA668433, [STDS0000058](https://db.cngb.org/stomics/mosta) [13], 10X Visium mouse brain (sagittal-anterior), 10X Visium mouse brain (sagittal-posterior), 10X Visium olfactory bulb, 10X Visium mouse kidney, and [ST mouse olfactory bulb](https://www.spatialresearch.org/resources-published-datasets/doi-10-1126science-aaf2403/) [15].
- Competing interests The authors declare no competing interests.
- Funding This work was supported by the National Key Research and Development Program of China (2023YFE0112300); National Natural Sciences Foundation of China (32261133526; 32270709; 32070677); the 151 talent project of Zhejiang Province (first level), the Science and Technology Innovation Leading Scientist (2022R52035); Jiangsu Collaborative Innovation Center for Modern Crop Production and Collaborative Innovation Center for Modern Crop Production cosponsored by province and ministry.
- Authors’ contributions S.L., Y.H., and M.C. conceptualized the study. S.L., Z.W., and Q.N. performed data analysis. S.L., M.C., Z.W., C.F., and S.Z. wrote the manuscript. All authors reviewed the manuscript.
- Acknowledgements

Editorial Policies for

Springer journals and proceedings: https://www.springer.com/gp/editorial-policies

Nature Portfolio journals: https://www.nature.com/nature-research/editorial-policies

*Scientific Reports*: https://www.nature.com/srep/journal-policies/editorial-policies

BMC journals: https://www.biomedcentral.com/getpublished/editorial-policies

---

**Fig. S1.**
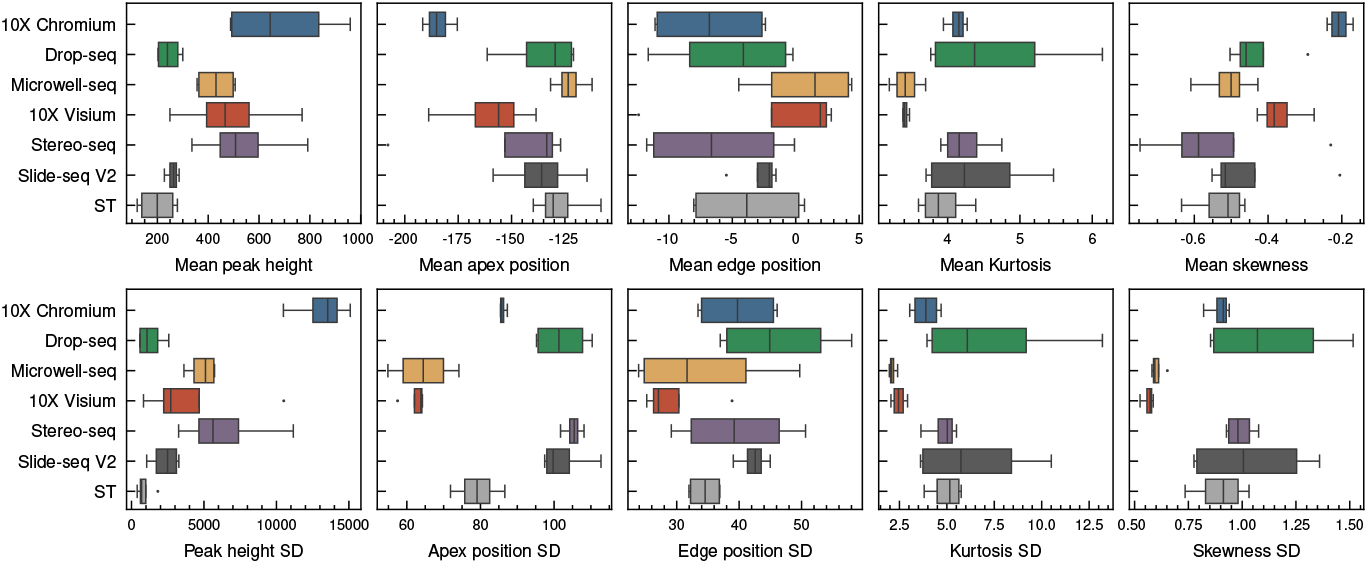
Comparison of average peak characteristics across different sequencing protocols. The peak characteristics include mean peak height, mean apex position, mean edge position, mean kurtosis, mean skewness, peak height standard deviation (SD), apex position SD, edge position SD, kurtosis SD, and skewness SD.

**Fig. S2.**
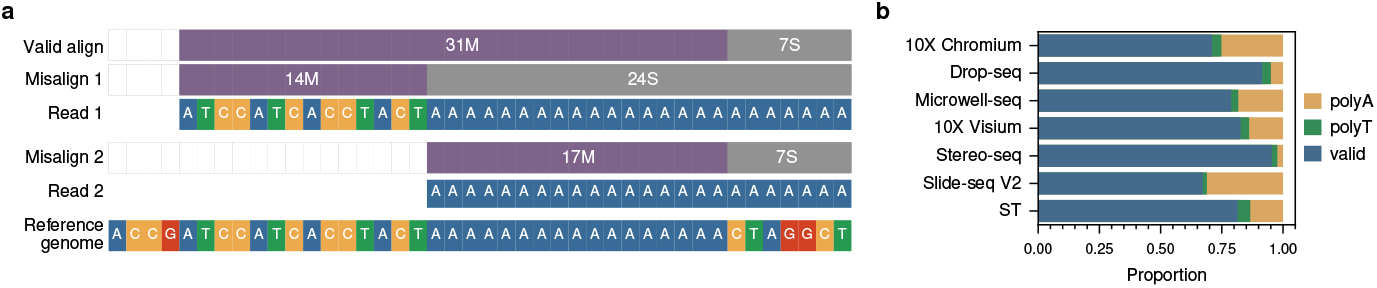
Artifacts in identifying polyA cleavage sites (pACSs). (**a**) Schematic representation of misalignment issues when aligning reads containing long polyA sequences to A-enriched regions of the reference genome. The alignment software may incorrectly determine the end position of the polyA sequence, leading to misalignment of the read. (**b**) Proportion of pACS reads containing polyA or polyT (defined as a stretch of at least 10 continuous A/T with at most one mismatch) in [-20,20] region across different sequencing protocols.

**Fig. S3.**
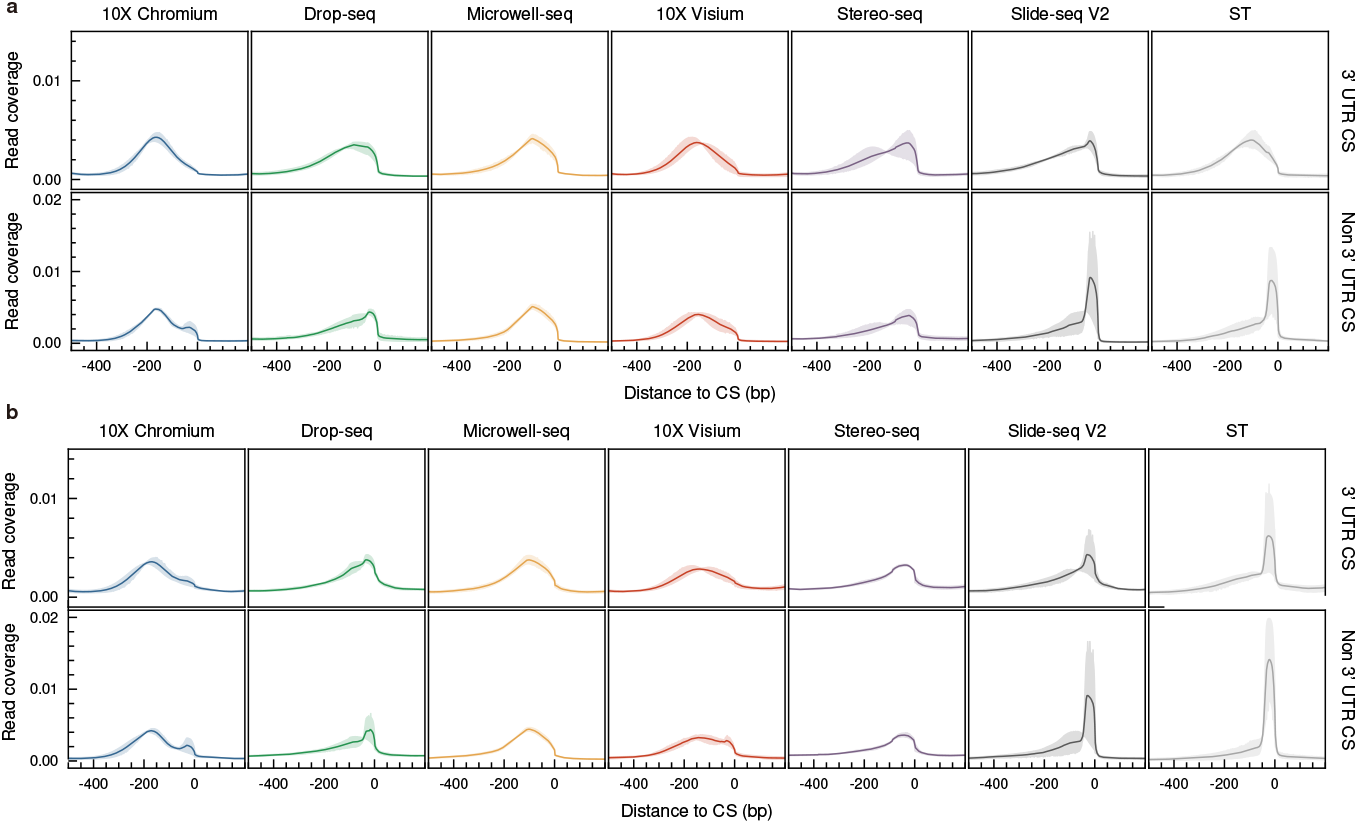
Read coverage profiles around polyA cleavage sites (pACSs) identified in different sequencing protocols. (**a**) Read coverage profiles around matched pACSs (i.e., pACSs that match annotated pA sites) across different sequencing protocols. (**b**) Read coverage profiles around unmatched pACSs (i.e., pACSs that do not match annotated pA sites) across different sequencing protocols.

**Fig. S4.**
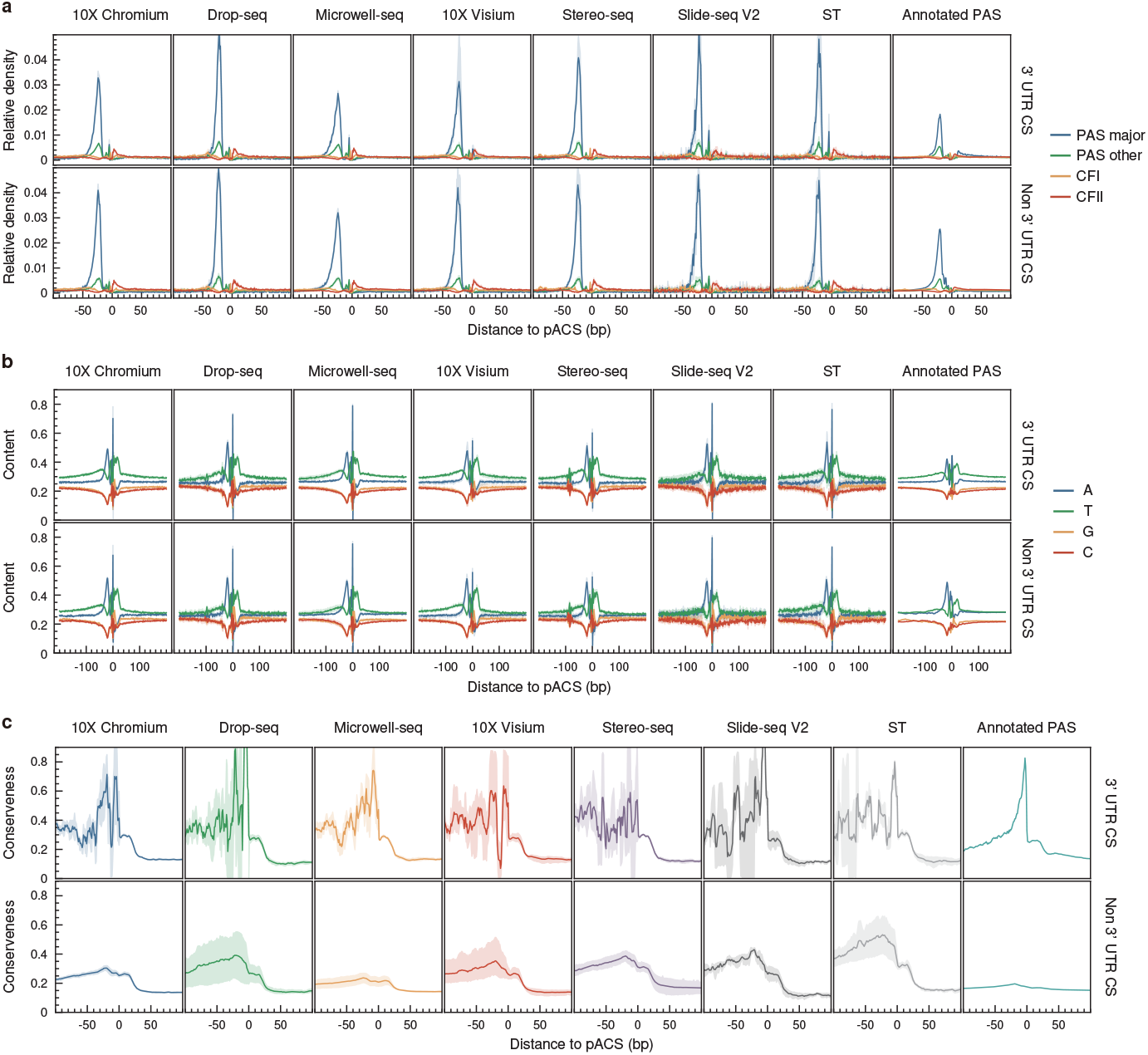
Sequence characteristics around matched polyA cleavage sites (pACSs) identified in different sequencing protocols and annotated pA sites. (**a**) Relative density of APA-related motifs (PAS major, PAS other, CFI, and CFII) around matched pACSs in 3’ UTR and non-3’ UTR regions. (**b**) Base composition around matched pACSs in 3’ UTR and non-3’ UTR regions. (**c**) Conservation profiles around matched pACSs in 3’ UTR and non-3’ UTR regions. Mouse phastCons60way.UCSC.mm10 conservation scores were extracted in 200 nt windows centered at pACS, with annotated coding sequence regions excluded.

**Fig. S5.**
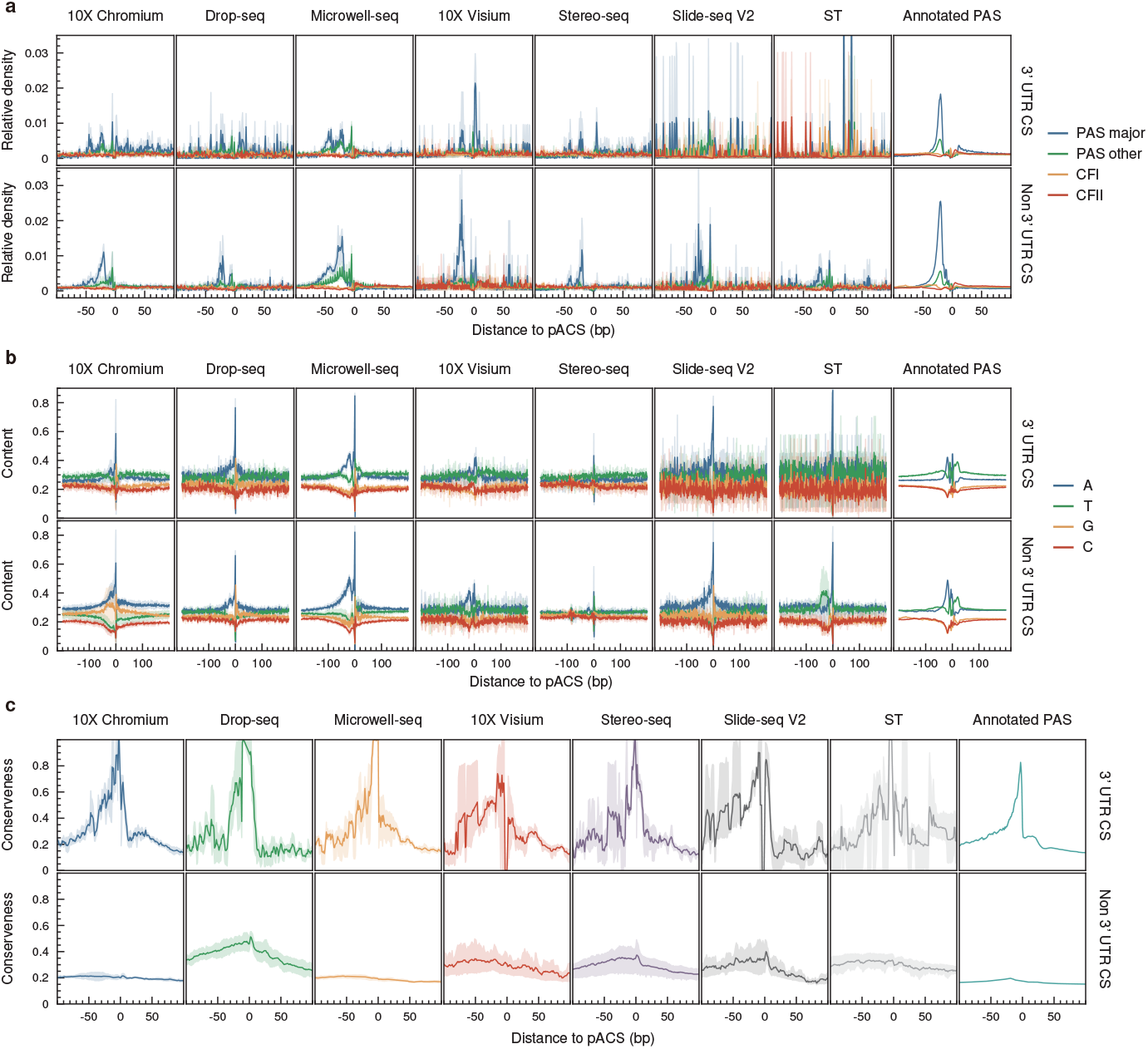
Sequence characteristics around unmatched polyA cleavage sites (pACSs) identified in different sequencing protocols and annotated pA sites. (**a**) Relative density of APA-related motifs (PAS major, PAS other, CFI, and CFII) around matched pACSs in 3’ UTR and non-3’ UTR regions. (**b**) Base composition around matched pACSs in 3’ UTR and non-3’ UTR regions. (**c**) Conservation profiles around matched pACSs in 3’ UTR and non-3’ UTR regions. Mouse phastCons60way conservation scores were extracted in 200 nt windows centered at pACS, with annotated coding sequence regions excluded.

**Fig. S6.**
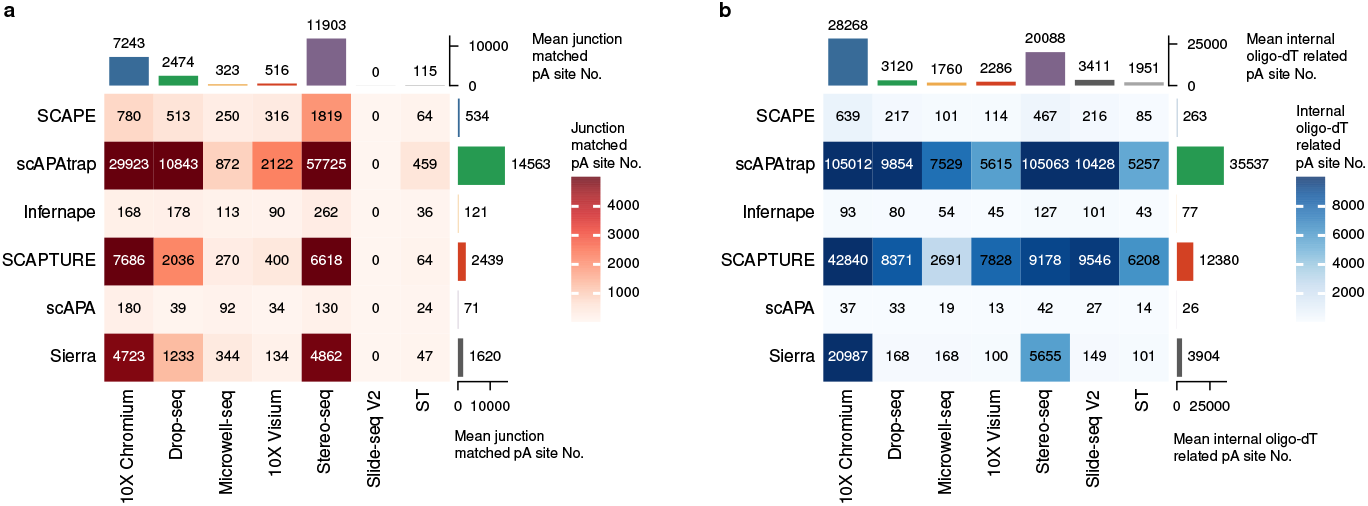
Evaluation of APA detection tools in filtering sequencing artifacts on simulated datasets.(**a**) Number of internal oligo-dT-related pA sites identified by each tool across different sequencing protocols. (**b**) Number of junction-matched pA sites identified by each tool across different sequencing protocols.

**Fig. S7.**
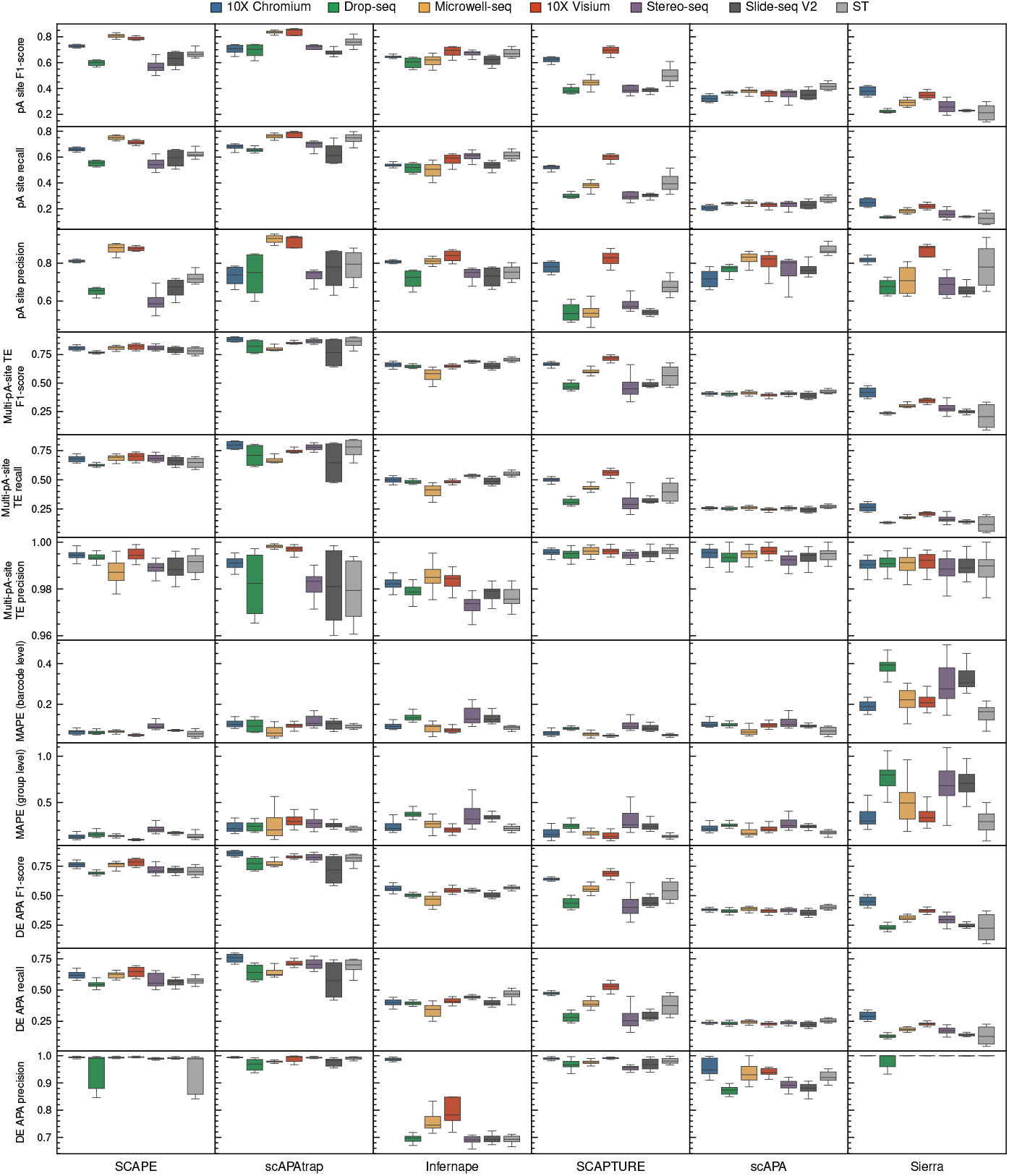
Performance evaluation of APA analysis tools (SCAPE, scAPAtrap, Infernape, SCAP-TURE, scAPA, and Sierra) on simulated datasets based on different sequencing protocols. The figure shows the F1-score, recall, and precision for pA site identification and multi-pA-site TE quantification, mean absolute percentage error (MAPE) for TE expression quantification at the barcode and group levels, and F1-score, recall, and precision for DE APA event detection.

**Fig. S8.**
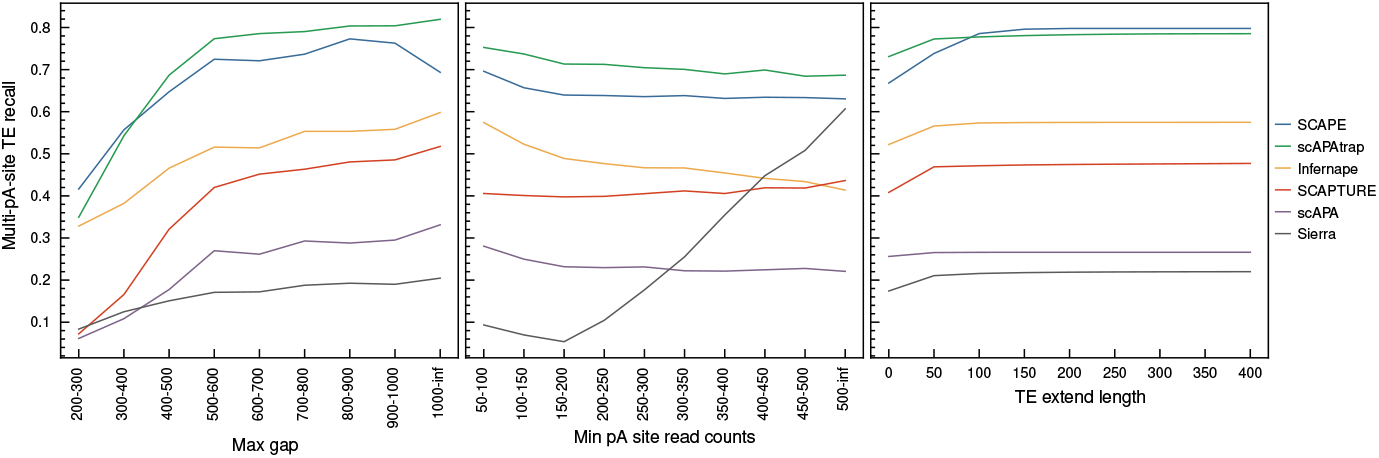
The effect of peak overlap (peak gap), weak signal (min peak size) and (TE extend length) on the recall rate of multi-pA-site TE. For each multi-pA-site TE unit, we ordered its ground truth pA sites according to their genomic positions and identified the pair of neighboring pA sites with the maximum gap. We then used the gap between these two pA sites and the read count of the smaller peak as the characteristic features representing the TE unit. For TE extend length, we extend te at different length to recovery pA sites that missed because of peak shift. The recall rate of multi-pA-site TE is shown with respect to the maximum gap between the selected pair of adjacent pA sites within the TE (left panel), the minimum read counts of pA sites of within the TE unit (middle panel), and the TE extension length (right panel).

**Fig. S9.**
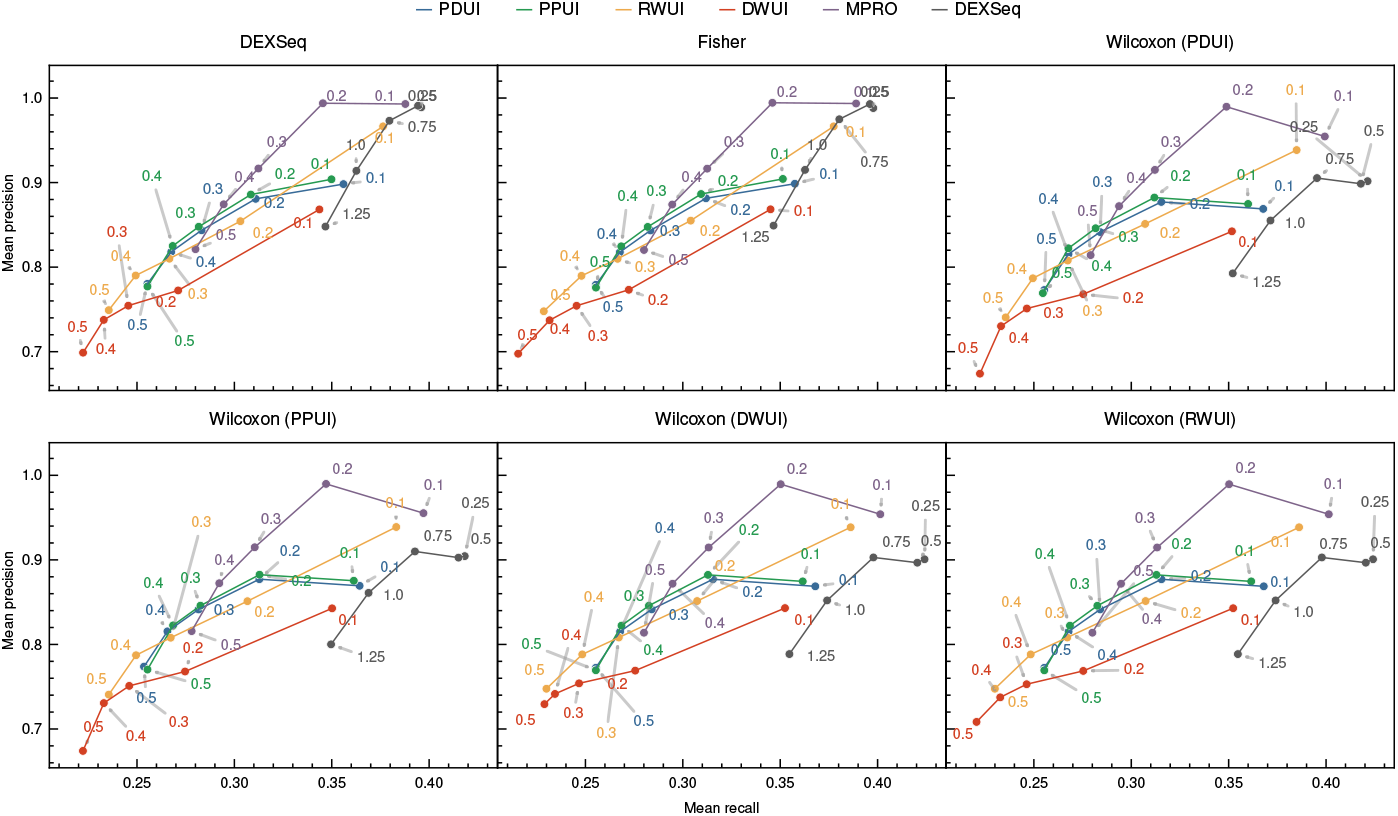
The effect of filtering thresholds on the performance of differentially expressed (DE) APA gene identification. The average precision and recall were shown for different combinations of filtering strategies (PDUI, PPUI, RWUI, DWUI, MPRO, and DEXSeq log2FC) and statistical tests (DEXSeq, Fisher’s exact test, and Wilcoxon rank-sum test) across an expanded range of threshold values.

